# Glucocorticoids reprogram human AML leukemic stem cells to promote elimination through differentiation and apoptosis

**DOI:** 10.64898/2026.09.09.750191

**Authors:** Isabella Angela Iasenza, Yi Ling Sun, LeRon Best, Hassan Dakik, Meaghan Boileau, Andrea L. Neumann, Patricia Arreba-Tutusaus, Manon Saby, Josée Hébert, Mark Minden, Luca Cavallone, Chris Williams, Bertrand Jean-Claude, Kolja Eppert

## Abstract

Acute myeloid leukemia (AML) is sustained by leukemic stem cells (LSCs) that can evade standard therapies and drive relapse. Targeting LSC-specific vulnerabilities is therefore essential for durable remission. Here we demonstrate that glucocorticoids (GCs) induce potent depletion of AML LSCs by promoting terminal differentiation and apoptosis. This effect is observable within 24 hours and is conserved across multiple LSC-enriched models and primary patient samples. Mechanistically, we establish that GC targeting of LSCs is mediated through the glucocorticoid receptor (NR3C1), with higher receptor binding affinity correlating with greater anti-LSC activity. We performed structure activity relationship (SAR) modeling of 24 corticosteroids and identified key features, including bulky D-ring substituents, associated with enhanced anti-LSC efficacy. Bulk and single-cell transcriptomic data revealed that GC treatment of LSCs suppresses NF-κB inflammatory signaling and disrupts stemness and quiescence programs while inducing transcriptional signatures associated with transient proliferation, metabolic stress, and terminal differentiation. Notably, GC sensitivity was associated with the expression of pre-existing inflammatory or extracellular matrix (ECM) signatures. Finally, we found that FLT3 ligand (FLT3L) is required for GC-induced proliferation of CD34- blasts but not for LSC depletion, suggesting that FLT3L levels may serve as a biomarker for blast expansion in patients receiving GC therapy. These findings support the clinical development of GC-based therapies in AML and provide mechanistic insights into how GCs target inflammatory and metabolic programs required for LSC survival.

## INTRODUCTION

AML is an aggressive hematologic malignancy marked by the uncontrolled proliferation of immature myeloblasts, leading to hematopoietic failure^1–3^. Despite high rates of initial remission, disease relapse remains the principal cause of treatment failure and mortality in AML. At the root of relapse is the LSC population, which possesses self-renewal capacity and can evade chemotherapy within protective bone marrow niches^3–11^. While induction regimens such as cytarabine (Ara-C) and anthracyclines can induce remission, they frequently fail to eliminate LSCs, necessitating therapies that specifically target this resistant compartment^12–17^.

LSCs exhibit distinct metabolic and inflammatory features compared with blasts and normal hematopoietic stem cells (HSCs), creating vulnerabilities that may be therapeutically exploited^12–17^. Pathways including NF-κB and CXCL12-CXCR4 are aberrantly activated in LSCs and reinforce quiescence and survival, while LSCs rely on oxidative phosphorylation (OXPHOS) for energy^17,18^. Disrupting these circuits may sensitize LSCs to therapy and promote their depletion. ^19^GCs are used in acute lymphoblastic leukemia (ALL) as standard of care to induce growth arrest and apoptosis in lymphoid cells but not used in AML treatment^20–23^. However, recent studies have renewed interest in the use of GCs in AML, identifying therapeutic activity in RUNX1 subtypes and NPM1-mutated AMLs^24–28^. Improved outcomes following GC treatment, particularly in combination with cytarabine, have been reported in patient-derived xenograft models and retrospective clinical study of patients with NPM1-mutated AML^26^. Additionally, AMLs that acquire resistance to cytarabine, venetoclax or FLT3 inhibitors can develop sensitivities to GCs, with several studies linking this response to heightened inflammatory transcriptional programs^26,28–31^. These findings support further investigation of GCs in AML but do not address their effects on LSCs, the population responsible for disease progression and relapse. In two previous studies, GCs displayed activity against LSCs or leukemia initiating cells *in vitro*, however, the nature of the LSC response to GC treatment and the molecular mechanisms underlying this activity remained unclear^26,32^.

In this study, we integrated transcriptomic, structural and functional analyses to investigate the effects of glucocorticoids on AML LSCs. We demonstrate that glucocorticoids act through the glucocorticoid receptor, NR3C1, to induce robust LSC depletion across genetically diverse cohort of primary AML samples. Mechanistically, GC treatment reprograms inflammatory and metabolic programs in LSCs and promotes terminal differentiation and apoptosis. We further show that this differentiation program can trigger a concurrent FLT3-dependent expansion of blasts, uncovering a previously unrecognized consequence of GC therapy. Together, these findings support the therapeutic potential of GCs in AML and identify a biomarker that may guide patient stratification.

## MATERIALS AND METHODS

### Culture and treatment of AML cell lines, patient samples, and LSC models

OCI-AML-8227, OCI-AML-20 (gift from Dr. Jean Wang, Princess Margaret Cancer Centre) and MUTZ-3 (DSMZ ACC 295) were cultured in cytokine-supplemented media and treated with glucocorticoids or vehicle control (0.01–0.02% DMSO (Fisher Scientific)) for four to six days. OCI-AML-20 cells were co-cultured with OP9 stromal cells (ATCC® CRL-2749™), while MUTZ-3 cells were maintained in 5637 bladder cell line conditioned medium. Cell viability and differentiation was assessed by flow cytometry using a LSR Fortessa with high-throughput sampler (HTS; BD Biosciences) after staining with antibodies to CD34 (APC), CD38 (PE), CD15 (FITC), CD14 (AlexaFluor® 700), CD45 (AlexaFluor® 700) (Biolegend) and SYTOX^TM^ Blue Dead Stain (Invitrogen^TM^). Combination assays were performed using MOM (0.01–1000 nM), cytarabine (10 nM, Tocris), or both, followed by analysis of differentiation and viability at four days (see supplemental methods).

Primary AML samples from the Ontario Cancer Institute (OCI1) (gifted by Dr. Mark Minden), Quebec Leukemia Cell Bank (BCLQ), and Nature Medicine cohort (OCI2) were cultured in OCI-AML-8227 media with the addition of 500 nM SR1 and UM729 (STEMCELL Technologies) treated with glucocorticoids for two to four days and analyzed by flow cytometry using lineage and maturation panels (see supplemental methods). OCI1 and OCI2 cohorts peripheral blood samples were collected from subjects with AML after obtaining informed consent according to the procedures approved by the Research Ethics Board of the University Health Network (Toronto). Sample 15H059 from the BCLQ was removed due to low cell viability prior to treatments.

### Apoptosis assay

Cell lines (Kasumi-1, MUTZ-3), AML patient samples (OCI1, BCLQ) and LSC models (OCI-AML-8227, OCI-AML-20) were plated in corresponding conditions and treated with MOM (Tocris) or DMSO (Fisher) for 24-96 hours (see supplemental methods). At each timepoint, cells were stained with CD34 (APC), CD38 (PE), CD15 (FITC), and CD14 (Alexa Fluor®700) (Biolegend), followed by Annexin V (Pacific Blue^TM^) and 7-AAD (Biolegend) to assess apoptosis. Cells were washed with PBS and Annexin V binding buffer (Biolegend) prior to incubation with viability dyes. Flow cytometry was performed using a BD LSRFortessa equipped with a high-throughput sampler.

### Structural-functional analysis (SAR) and molecular modeling of corticosteroids

Molecular structures of corticosteroids were obtained from PubChem and grouped based on functional group composition using the Coopman classification system (Groups A, B, C, D1, D2). Key structural features associated with anti-LSC activity were identified by comparing effective and ineffective compounds within each class. To model ligand-receptor interactions, Protein Data Bank (PDB) files for MOM furoate (4P6W), triamcinolone acetonide (5UFS), dexamethasone (1M2Z), and hydrocortisone (6NWL) were analyzed using Molecular Operating Environment (MOE) software. Structures were aligned to assess conserved features, steroid core positioning, and protein helix alignment (RMSD). Electrostatic maps and hydrophobic surface modeling were used to evaluate binding pocket interactions, and ligand activity was correlated with percent survival of CD34+CD38- cells at 15 nM.

### Antagonism experiment with glucocorticoids and RU486

OCI-AML-8227 cells were plated and treated with the glucocorticoid receptor antagonist RU486 (Tocris) at various concentrations, either alone or in combination with 1 nM MOM (Tocris) or 20 nM DEX (Sigma). After 4 days, cells were collected, stained with antibodies against CD34 (APC), CD38 (PE), CD15 (FITC) (Biolegend), and SYTOX (Life Technologies), and analyzed by flow cytometry using an LSR Fortessa with high-throughput sampler (HTS). Data were normalized to the glucocorticoid-only condition, and statistical significance was assessed by unpaired t-test using GraphPad Prism v10.

### Cytokine experiments

For the FLT3L withdrawal assay, bulk and FACS-sorted OCI-AML-8227 cells were cultured in the absence of one growth factor at a time (IL3, SCF, IL6, TPO, GCSF, FLT3L) and treated with increasing doses of MOM.

For the FACS-sorted experiments, OCI-AML-8227 was enriched using EasySep™ CD34 Positive Selection kit II (STEMCELL Technologies, #17856), expanded for four weeks, and sorted into CD34+ (LSPCs) and CD34- (blast) fractions using a BD FACSAria™ Fusion sorter. Cells (10,000/well) were plated with or without FLT3L and treated with MOM or DMSO for five days. Cells were stained with CD34 (APC), CD38 (PE), CD15 (FITC), and SYTOX (Life Technologies), and analyzed by flow cytometry. LC₅₀ values were calculated using GraphPad Prism (nonlinear regression), and statistical comparisons were performed using paired t-tests.

### RNA-sequencing

RNA integrity was confirmed on an Agilent 2100 Bioanalyzer. Libraries were prepared with the Illumina Stranded mRNA Prep Ligation kit, quantified with the KAPA Library Quantification kit, and sequenced using an Illumina NovaSeq 6000 S4 platform (paired-end 2^100 bp). Fastq files were processed on Compute Canada clusters using the RNA-seq module of Genpipes (v4.4.2) with default parameters^33^. Briefly, raw reads were trimmed with Trimmomatic ^34^ (v0.36) to remove Illumina adapters and sequencing-primer associated reads. High-quality reads were then aligned to the hg38 (GRCh38) human genome build using 2-pass STAR^35^ (v2.7.8a). PCR duplicate reads were collapsed by Picard (v2.9), and gene annotation was based on the Ensembl database (human release 104). Differential gene expression was done in R environment using the edgeR package, and genes with BH-adjusted p<0.05 and |log?FC | > 1 or 1.5 were considered significant. Gene set enrichment analysis (GSEA) and VennDiagram were used to assess pathway enrichment and gene overlaps. Full protocol and flow cytometry validation are described in the Supplemental Methods.

### Single-cell RNA sequencing of enriched CD34+ LSPCs OCI-AML-8227 cells

Enriched CD34+ LSPCs OCI-AML-8227 cells were treated with 0.5 nM MOM or DMSO (0.05%) for 24 hours, washed, and resuspended in PBS + 0.04% BSA for scRNA-seq. Single-cell libraries were prepared using the 10x Genomics Chromium X system and Single Cell 3’ v3.1 reagents, targeting ∼10,000 cells per condition. Libraries were sequenced on an Illumina NovaSeq 6000 (28 bp Read1, 150 bp Read2, dual 8 bp indexes). Raw BCL files were demultiplexed with bcl2fastq, aligned to GRCh38, and processed using cellranger count v7.1.0 (10x Genomics). Count matrices were analyzed using Seurat, including clustering, dimensionality reduction, and cell type annotation (details in Supplemental Methods). Single-sample GSEA (ssGSEA) was performed using GSVA to assess pathway-level changes across annotated cell types.

### *Ex vivo* xenograft assays

Mouse experiments were performed according to protocols approved by McGill University and its Affiliated Hospital’s Research Institutes. OCI-AML-8227 cells were plated in corresponding media and treated with 20 nM MOM (Tocris) or 0.04% DMSO (Fisher Scientific) for seven days. Cells were then washed, counted, and stained with a panel including CD45 (Alex Fluor®700), CD19 (BV711), CD33 (APC), CD11b (BV650), CD15 (FITC), CD34 (APC Cy7), CD38 (PE) (Biolegend) and SYTOX (Pacific Blue™) (Life Technologies). Cells (25 ^L total volume, 1/4 bulk population) were injected intrafemorally into sublethally irradiated NSG-S mice (2.1 Gy, X-RAD SmART Irradiator; Precision X-Ray, Inc). Mice were sacrificed after 12 weeks, and human engraftment (CD45⁺) was evaluated in injected femur, contralateral femur, and spleen by flow cytometry.

## RESULTS

### Glucocorticoids deplete LSCs through differentiation without apoptosis within 24 hours

It has been previously demonstrated that GC treatment depletes LSC-enriched cell populations with a concurrent increase in differentiated cells after six days in the OCI-AML-8227 model, suggesting LSC differentiation through GCs^32^. However, it remained unclear whether the loss of LSCs was driven by apoptosis or by differentiation into downstream progenitor populations. To investigate the kinetics and mechanism of this response, we compared four glucocorticoids, mometasone (MOM), halcinonide, budesonide and dexamethasone, in a six-day dose response assay. MOM consistently produced the most robust and reproducible depletion of the LSC-enriched CD34+CD38- population and was therefore selected for subsequent mechanistic studies (Supp. Figure 1A, B).

To characterize the early response to GC treatment, we performed a four-day time course and apoptosis assay in OCI-AML-8227, assessing CD34+CD38- LSCs, CD34+ leukemic stem and progenitor cells (LSPCs) in combination, and CD34- blast populations following MOM treatment (Figure 1A, B). In human AML samples, LSC-enriched populations (CD34+CD38-) can differentiate to leukemic progenitor cells (LPCs) (CD34+CD38+), which in turn can differentiate to leukemic blasts (CD34-). A significant reduction in the LSC-enriched fraction was observed as early as 24 hours, with a corresponding increase in CD34- blast cells (Figure 1A). Annexin V staining revealed no increase in apoptosis across any populations or timepoint following treatment, suggesting that LSC depletion occurs independently of cell death and is consistent with differentiation (Figure 1B).

**Figure 1:**
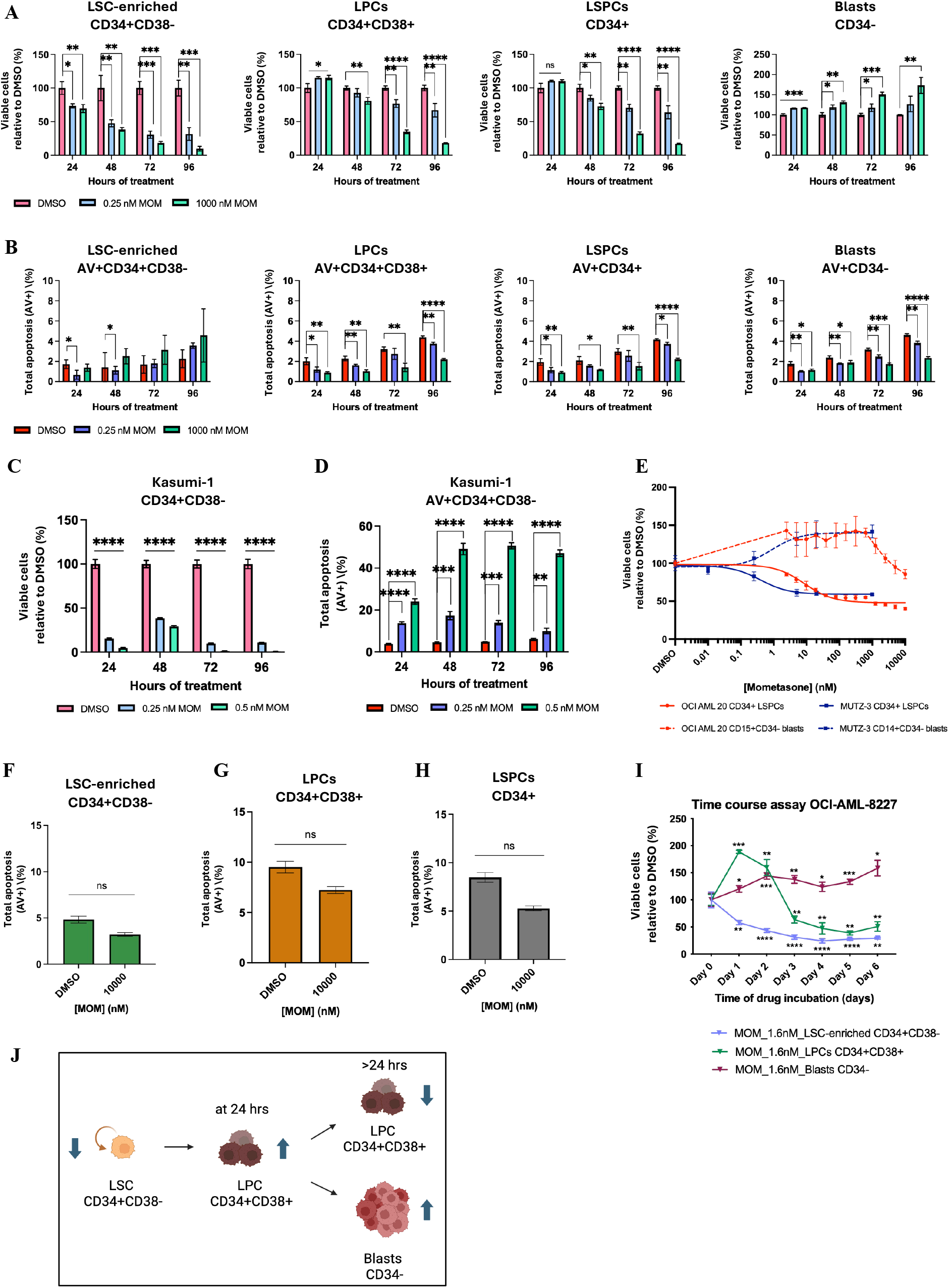
Glucocorticoids deplete LSCs through terminal differentiation as early as 24 hours and without apoptosis across AML LSC models. **(A)** Time course analysis of OCI-AML-8227 cells treated with MOM (MOM, 0.25 and 1,000 nM) for 24 to 96 hours, assessing changes in CD34+ and CD34- populations. Cell viability relative to DMSO control is shown for CD34- blasts, CD34+ leukemic stem and progenitor population (LSPCs), CD34+CD38+ leukemic progenitors (LPCs) and CD34+CD38- (LSC-enriched population). P value calculated using unpaired student t-test. Mean ± s.d. Representative one of *n* = 3 biological replicates. **(B)** Apoptosis from time course in **(A)** (24 to 96 hrs) of OCI-AML-8227 treated with MOM. Annexin V (AV) and 7-AAD were used to assess apoptosis. Total AV+ cells in each cell fraction are shown. P value calculated using unpaired student t-test. Mean ± s.d. Representative one of *n* = 3 biological replicates. **(C-D)** Kasumi-1 time course treated with MOM (0.25 and 0.5 nM) for 24 to 96 hrs. Cell viability **(C)** and apoptosis **(D)** is shown for CD34+CD38- cells. Apoptosis was determined using Annexin V and 7-AAD. Total AV+ cells in each cell fraction are shown. P value calculated using unpaired student t-test. Mean ± s.d. Representative one of *n* = 3 biological replicates. **(E)** Dose response treatment of LSC-models OCI-AML-20 and MUTZ-3 with various concentrations of MOM for five days. CD34+ LSPCs and blast populations of both are shown (OCI-AML-20: CD15+CD34- blasts, and MUTZ-3: CD14+CD34- blasts). P values calculated using unpaired student t-test. Mean ± s.d. Representative one of *n* = 3 biological replicates. **(F-H)** Bar plot of total apoptosis (%AV+) in CD34+ LSPCs, CD34+CD38+ LPCs and CD34+CD38- LSC-enriched fraction of OCI-AML-20 cells at highest dose of MOM (10,000 nM) for five days. P value calculated using unpaired student t-test. Mean ± s.d. Representative one of *n* = 3 biological replicates. **(I)** Six-day time course assay of OCI-AML-8227 treated with 1.6 nM of MOM. Viable CD34- blasts, CD34+CD38+ LPCs and CD34+CD38- LSC fraction is shown relative to DMSO. Mean ± s.d. Representative one of *n* = 2 biological replicates. **(J)** Proposed LSC differentiation model with glucocorticoid treatment. MOM targets CD34+CD38- cells and drives them to differentiate to CD34+CD38+ LPCs as early as 24 hours. These further differentiate into CD34- blasts. LPC further decrease due to differentiation after 24 hours of treatment. Apoptosis does not occur. * p < 0.05, ** p < 0.01, *** p < 0.001 and **** p < 0.0001.

To verify that our assay could detect GC-induced apoptosis and to determine whether differentiation represented a general response to GC treatment, we included Kasumi-1 cells as a control. This *RUNX1*-rearranged AML cell line has been previously reported to undergo apoptosis in response to GC treatment^24^. As expected, Kasumi-1 cells showed a strong apoptotic response (>50% Annexin V⁺) without differentiation at all timepoints post-treatment, in contrast to the OCI-AML-8227 model (Figure 1C, D, Supp. Figure 1C, D). Furthermore, to determine whether GC-induced LSC differentiation was generalizable beyond OCI-AML-8227, we tested MOM in two additional LSC-containing AML models of adverse-risk *EVI1*-driven AML: OCI-AML-20 and MUTZ-3 (Figure 1E). Both models harbor CD34+ LSPC populations and are amenable to *in vitro* differentiation (Supp. Figure 1E, F). MOM significantly reduced the CD34+ LSPC populations in both models (e.g., OCI-AML-20: 100% vs. 39.8%, p < 0.0001; MUTZ-3: 100% vs. 59%, p = 0.0003), while inducing expansion to CD15+CD34- blasts in OCI-AML-20 (p = 0.0003) and CD14+CD34- blasts in MUTZ-3 (p = 0.005) (Figure 1E). Apoptosis levels remained low, and stromal viability was not impaired, supporting a shared mechanism of differentiation of LSCs across models (Figure 1F-H, Supp. Figure 1G).

To further define the differentiation trajectory, we performed a six-day kinetic assay evaluating changes across all cell populations in OCI-AML-8227: LSC-enriched fraction, leukemic progenitors, and blasts (Figure 1I, Supp. Figure 2A-D). The LSC-enriched population rapidly declined post-treatment, whereas the leukemic progenitors transiently accumulated and peaked at day one before subsequently decreasing, suggesting an intermediate differentiation state (Figure 1I, Supp. Figure 2A-D). These findings support a hierarchical differentiation process in which GC treated LSCs pass through a progenitor intermediate before generating blast populations (Figure 1J).

Together, these results demonstrate that glucocorticoids deplete AML LSC-enriched populations through a differentiation associated process rather than apoptosis across multiple LSC-containing models (Figure 1J). These findings support the therapeutic potential of GCs as differentiation inducing agents in AML and provide a framework for investigating the molecular pathways governing GC-mediated LSC depletion.

### Glucocorticoids target LSPCs across genetically diverse primary AML samples

To evaluate the sensitivity and type of response to glucocorticoids in primary human AML, we treated 30 patient samples with MOM across three cohorts: Ontario Cancer Institute 1 (OCI1; *n* = 15), Ontario Cancer Institute 2 (OCI2; *n* = 6), and Leukemia Cell Bank of Quebec (BCLQ; *n* = 9) (Supp. Tables S1-S3). As LSCs can often be observed in both the CD34+ CD38- and CD34+ CD38+ fractions of primary AML samples, we assayed the CD34+ LSPC population in these cohorts^36^. From the nine primary patient samples from the BCLQ, MOM treatment depleted CD34+ LSPCs in all samples, confirming glucocorticoid sensitivity across genetically diverse AMLs (Figure 2A, B Supp. Table S4). These primary AMLs exhibited heterogenous cellular responses to GC treatment: 33% of samples had their CD34+ LSPCs depleted through apoptosis only whereas 67% of the samples depleted the CD34+ LSPCs through differentiation in the absence of apoptosis (Figure 2A, B). These findings indicate that GCs can eliminate LSPCs through distinct cellular responses.

**Figure 2:**
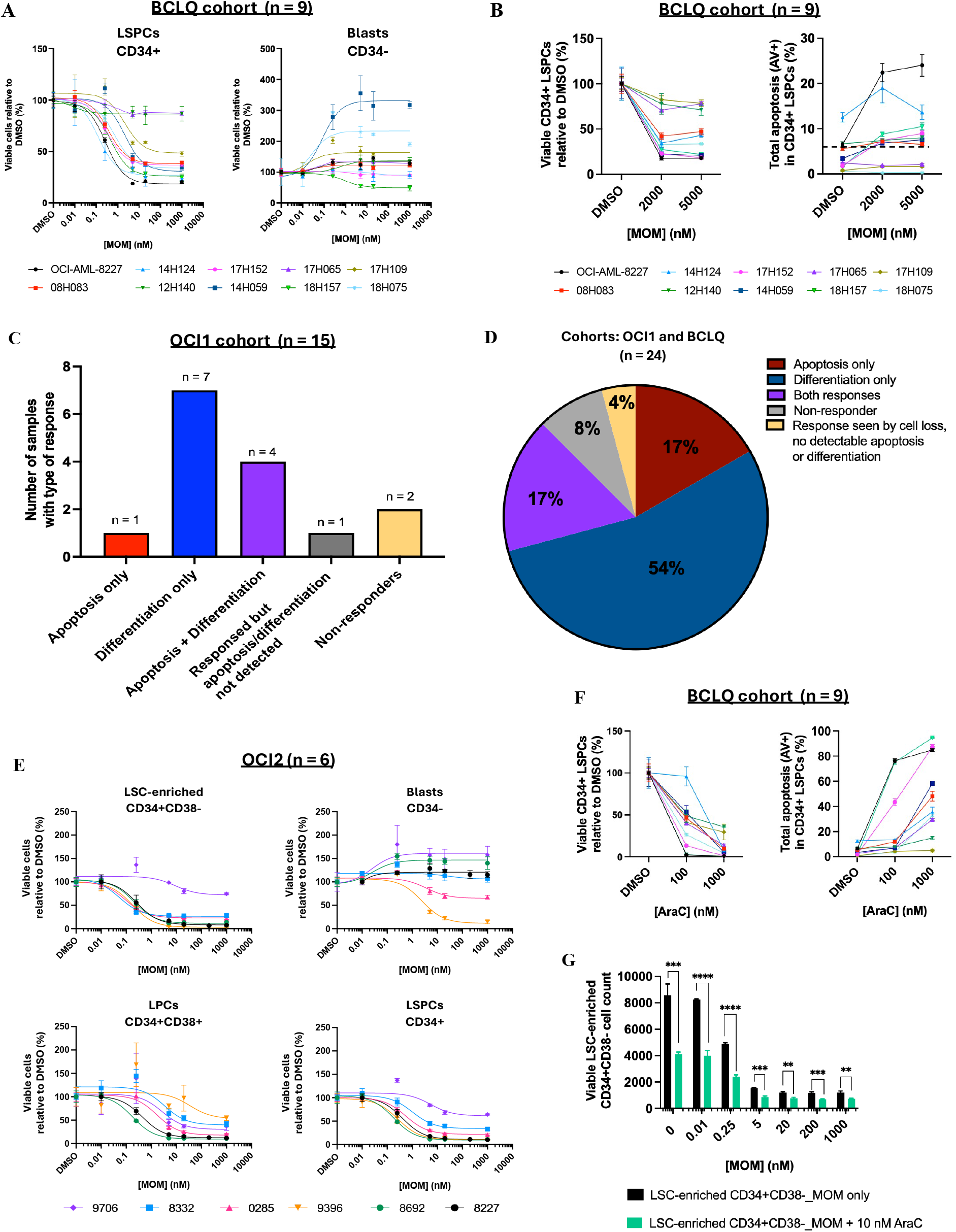
MOM targets primary AML LSCs and enhances cytarabine efficacy. **(A)** Dose response of BCLQ cohort (*n* = 9) treated with MOM for five days. MOM targets CD34+ LSPCs and induces differentiation of some samples to CD34- blasts. Viable cells relative to DMSO control is shown. Mean ± s.d. OCI-AML-8227 was used as a control. **(B)** Viable cells relative to DMSO for (left) CD34+ LSPCs from BCLQ cohort ( *n* = 9) treated with 2,000 and 5,000 nM of MOM for three days. (Right) Apoptosis was measured using Annexin V and 7-AAD. MOM induces apoptosis in some samples. Dotted line indicates apoptosis cut-off (>5% AV+ and >5% increase from DMSO control). **(C)** Bar plot summarizing the types of responses to MOM from OCI1 cohort (*n* = 15) treated for three days. Majority of samples had the LSPC CD34+ fraction depleted through differentiation only. Number of samples that have corresponding response is indicated above bars. *N* = 1 biological replicate. **(D)** Response classification of AML patient samples from 24 samples with phenotypic data treated with MOM. The majority of the samples (54%) responded with LSC- induced differentiation, while 17% responded with apoptosis only. A proportion of samples (17%) had both types of responses, and a few (8%) did not respond to treatment. Responded but non- detected refers to a depletion in cell count of the corresponding population but differentiation and/or apoptosis was not observed (4%). Representative one of *n* = 1 biological replicates for all patient cohorts. **(E)** Dose response of OCI2 cohort AML samples (*n* = 6, confirmed LSC-enriched fractions (Supp. Table S3)) treated with MOM for four days. LSC-enriched fractions are depleted, and some samples respond through differentiation. **(F)** Viable cells relative to DMSO for (left) CD34+ LSPCs from BCLQ cohort ( *n* = 9) treated with 100 and 1,000 nM of cytarabine (Ara-C) for three days. (Right) Apoptosis was measured using Annexin V and 7-AAD. Total apoptosis in CD34+ LSPCs is shown. Ara-C induces apoptosis in BCLQ cohort but does not deplete LSPCs through differentiation. OCI-AML-8227 was used as a control. Mean ± s.d. **(G)** Combination of MOM and 10 nM Ara-C for four days in OCI-AML-8227 CD34+CD38- LSC-enriched fraction. Unpaired t-test, * p < 0.05, ** p < 0.01, *** p < 0.001 and **** p < 0.0001. Mean ± s.d. Representative one of *n* = 2 biological replicates.

In the OCI1 cohort, 87% of the samples responded to MOM with a depletion of the CD34+ LSPC population enriched for both LSCs and progenitors (Figure 2C, Supp. Table S4). The responses were heterogenous among samples: seven samples underwent differentiation associated depletion, similar to OCI-AML-8227, characterized by the loss of CD34+ LSCPs with a corresponding expansion of CD34- blasts in the absence of apoptosis, whereas only one responded through apoptosis alone and four samples exhibited both apoptosis and differentiation. Furthermore, one sample displayed depletion of the CD34+ LSPCs without apoptosis or differentiation detected and two samples were non-responders (Figure 2C, Supp. Table S4). Like the OCI-AML-8227 model, a substantial subset of primary AMLs CD34+ LSPCs underwent depletion through differentiation. Overall, in both OCI1 and BCLQ cohorts, MOM depleted the LSPCs in 92% of the 24 tested AML samples, with 54% responding through differentiation with an absence of apoptosis, 17% through apoptosis alone, 17% through a combination of both mechanisms, and 4% with depletion but neither mechanism detected (Figure 2D).

To determine whether GCs target functionally validated LSC populations rather than the broader CD34+ LSPC population, we analyzed the third cohort (OCI2) of six primary AML samples that have been confirmed to harbor leukemia-initiating cells in their CD34+CD38- fractions via xenotransplantation, including OCI-AML-8227^36,37^. MOM targeted the CD34+CD38- LSC-enriched populations in 83% of samples, with LC₅₀ values in the low nanomolar range, and half exhibited a corresponding increase in CD34- blasts, consistent with LSC differentiation (Figure 2E). These findings demonstrate that GC activity extends to functionally validated AML LSC populations.

Together, these results demonstrate the consistent sensitivity of LSPCs to GC treatment across primary AML samples and that depletion can occur through either differentiation or apoptosis.

### Glucocorticoids exhibit activity distinct from cytarabine and enhance LSC depletion upon combination treatment

Given the heterogenous responses elicited by MOM across primary AML samples, we next compared its activity with cytarabine (Ara-C), a standard chemotherapeutic agent used in AML, alone and in combination.

Treatment of primary AMLs with Ara-C depleted viable CD34+ LSPCs across BCLQ samples, primarily through apoptosis (Figure 2F). In contrast, MOM depleted CD34+ LSPCs but also promoted expansion of the CD34- blast population in several samples, consistent with the differentiation associated responses observed in OCI-AML-8227 and a subset of primary samples from previous cohorts tested (Figure 2A-B). These findings suggest that MOM can induce cellular responses distinct from those induced by conventional cytotoxic chemotherapy.

Due to MOM inducing a differentiation associated response in the LSC-enriched populations of OCI-AML-8227 and in a substantial subset of primary AMLs, we next investigated whether differentiation inducing GC-treatment could complement Ara-C mediated cytotoxicity. Co-treatment significantly enhanced depletion of the CD34+CD38- LSC-enriched population compared to either treatment alone (Figure 2G). For example, combining 10 nM Ara-C with increasing doses of MOM results in a >25% additional depletion of CD34+CD38- LSC-enriched cells relative to Ara-C alone (48% vs. 28% remaining viable cells with 0.25 nM MOM + Ara-C, p< 0.001; Figure 2G), demonstrating cooperativity between the two drugs. Together, these findings demonstrate that MOM can enhance Ara-C mediated depletion of the LSC-enriched populations in the OCI-AML-8227 model due to distinct responses.

### Glucocorticoid response is not associated with specific mutations but correlates with pre-treatment inflammatory and ECM transcriptional programs

To determine whether specific genetic alterations might influence the sensitivity of LSCs to GC treatment in human AML, we genotyped for common AML-associated mutations. Genotyping across two cohorts, OCI1 and OCI2, confirmed that glucocorticoid response in LSCs did not associate with specific mutations or cytogenetic features (Supp. Table S5). In addition, the level of expression of the glucocorticoid receptor (GR), *NR3C1*, was not linked to response (Supp. Figure 3A).

These findings prompted us to investigate whether non-genetic features, such as transcriptional programs present prior to treatment, might underlie sensitivity to GCs. We performed RNA-seq on 21 primary AML samples from the cohorts prior to treatment with known phenotypic responses to MOM. Seven samples lacked apoptosis information or were non-responders and were therefore excluded from the apoptotic analyses. In the samples that responded with apoptosis upon treatment, several genes were significantly upregulated prior to treatment compared to samples without apoptosis, including *COL6A2*, *TPSB2*, and *NEO1*, which are involved in extracellular matrix (ECM) remodeling and immune signaling (Figure 3A, Supp. Figure 3B). Gene set enrichment analysis (GSEA) of these samples identified consistent enrichment of ECM and immune-related pathways such as collagen trimerization and complement activation prior to treatment (Figure 3B, Supp. Table S6). In contrast, AML samples that responded through GC-induced differentiation upon treatment expressed higher levels of inflammatory genes prior to treatment such as *SOCS3*, *IFI44L*, and *NFKBIZ* compared to samples without differentiation (Figure 3C, Supp. Figure 3C). These samples showed enrichment of immune and inflammatory pathways, including scavenger receptor activity and NF-κB–mediated signaling, while non-responders had few upregulated programs prior to treatment (Figure 3D, Supp. Table S7). These data suggest that inflammatory signatures are associated with differentiation, whereas ECM programs correlate with apoptosis prior to treatment.

**Figure 3:**
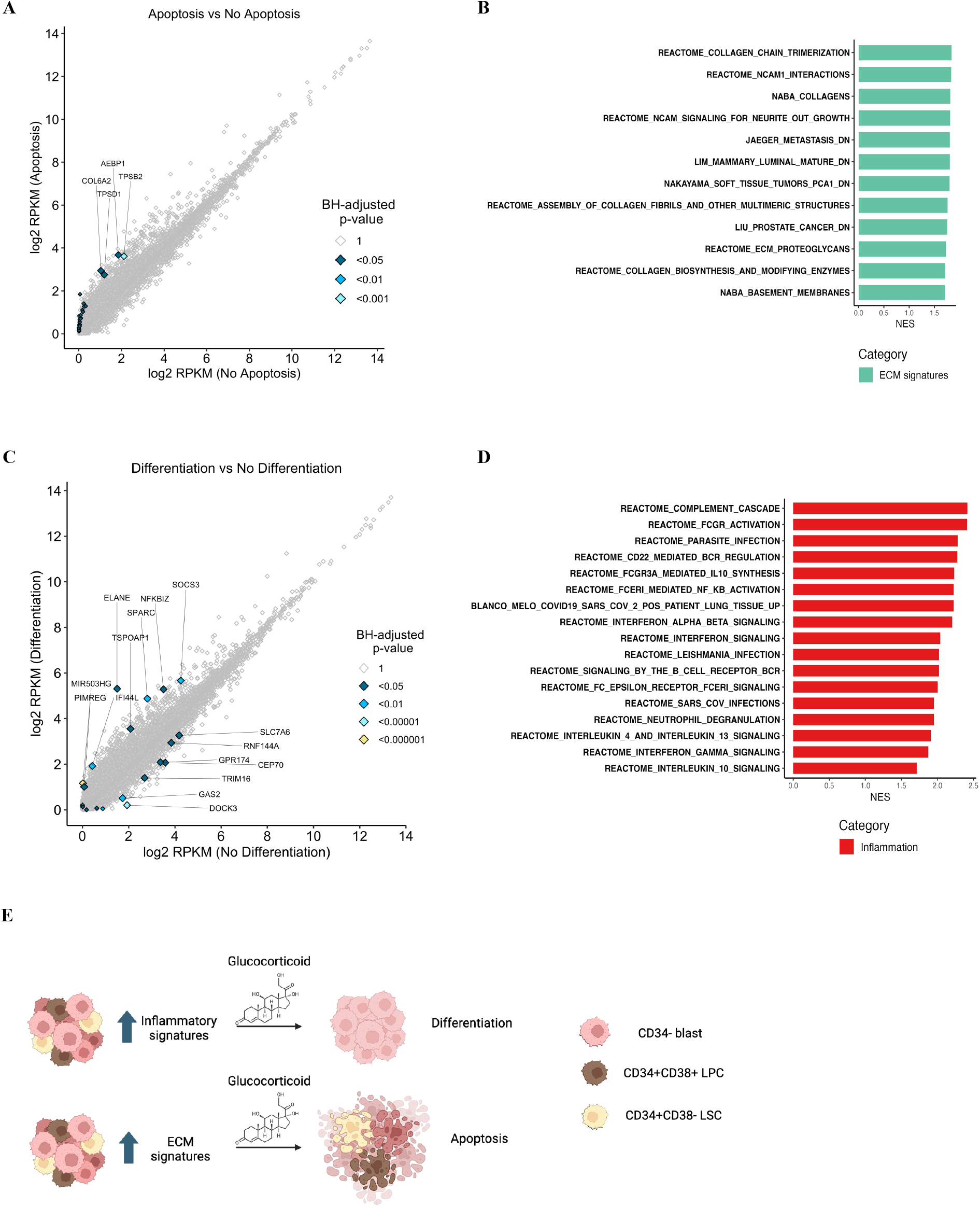
Glucocorticoid response is independent of mutations but correlates with inflammatory and ECM gene signatures. **(A)** Differentially expressed genes (log_2_RPKM) in pre-treated AML apoptotic responders compared to those that do not undergo apoptosis (*n* = 14 samples used for sequencing, seven samples did not have apoptotic information or were non- responders). Genes are color-coded based on BH-adjusted p-value. Significant genes are selected based on BH-adjusted p < 0.05, log_2_FC cut-off = 1; min average log_2_RPKM per group = 2. See FC table – Apoptosis vs NoApoptosis. Highly expressed genes were found to be involved in extracellular matrix (ECM) remodeling. **(B)** GSEA of apoptotic responders shows enrichment of ECM pathways. NES: normalized enrichment score. Shown are gene sets with p < 0.05 and FDR < 0.25. **(C)** Differentially expressed genes (log_2_RPKM) in pre-treated AML differentiation responders compared to those that did not undergo differentiation (*n* = 21 samples used for sequencing). Genes are color-coded based on BH-adjusted p-value. Significant genes are selected based on BH-adjusted p < 0.05, log_2_FC cut-off = 1.5; min average log_2_RPKM per group = 1. See FC table – Diff vs No Diff. Highly expressed genes were found to be involved in inflammatory response. **(D)** GSEA of differentiation responders shows enrichment of NF-κB and scavenger pathways. Gene sets shown have p < 0.05 and FDR < 0.25. NES: Normalized enrichment score. **(E)** Proposed model of LSC response to glucocorticoids. LSCs, when driven out of quiescence into LPCs, must either differentiate or undergo apoptosis. Genetic signatures pre-treatment associated with ECM remodeling favor apoptosis while inflammatory signatures favour differentiation.

Together, these results suggest that the type of response LSCs undergo with GC treatment is not dictated by specific mutations, but rather by baseline transcriptional programs prior to treatment: AMLs enriched in inflammatory signatures preferentially may undergo differentiation, whereas those with ECM-like features may be more prone to apoptotic response (Figure 3E).

### Glucocorticoids require NR3C1 engagement and specific structural features to promote LSC depletion

Glucocorticoids act through the glucocorticoid receptor (NR3C1) to regulate downstream transcriptional programs. To determine whether GR signaling is required for GC-mediated LSC depletion, we performed GR antagonism studies in OCI-AML-8227 cells using RU486. Co- treatment with RU486 blocked the effects of MOM, preventing depletion of the CD34⁺CD38⁻ LSC-enriched fraction and reducing the expansion of CD34- blasts (Figure 4A, B). Similar results were observed with dexamethasone (Supp. Figure 4A-D), indicating that GR activation is required for GC-induced differentiation and LSC depletion in the OCI-AML-8227 model.

**Figure 4:**
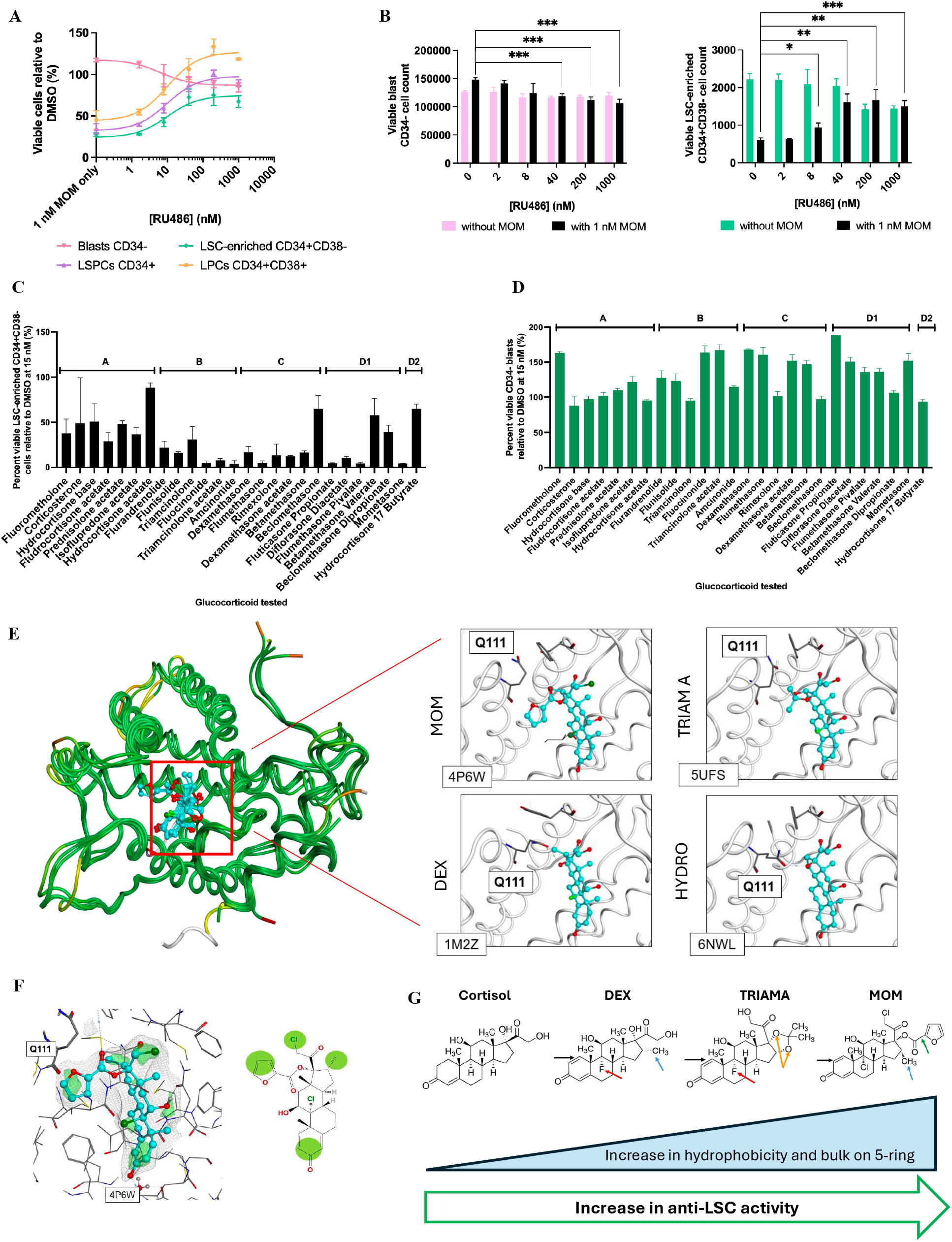
Glucocorticoids bind NR3C1 to induce LSC differentiation, with binding affinity correlating with anti-LSC potency. **(A)** RU486 blocks MOM-induced LSC differentiation. OCI- AML-8227 cells were treated with 1 nM MOM with or without the glucocorticoid receptor (NR3C1) antagonist RU486 for six days. RU486 prevented depletion of the CD34+ LSPCs, CD34+CD38+ LPCs and CD34+CD38- LSC-enriched populations and CD34- blast expansion, confirming that NR3C1 signaling is required for MOM-induced differentiation. Viable cells relative to DMSO is shown. Mean ± s.d. Representative one of *n* = 3 biological replicates. **(B)** Viable cell count for CD34- blasts (left) and CD34+CD38- LSC-enriched fraction (right) for OCI- AML-8227 treated with 1 nM MOM ± RU486. Starting dose of 0 nM for colored bars indicate DMSO only without RU486, for black bar indicates 1 nM of MOM is present. As concentrations increase, RU486 is added to both conditions. Unpaired t-test, * p < 0.05, ** p < 0.01, *** p < 0.001 and **** p < 0.0001. Mean ± s.d. Representative one of *n* = 3 biological replicates. **(C-D)** Twenty- four corticosteroids spanning 5 structural subtypes (A,B,C,D1 and D2) were screened in OCI- AML-8227 cells for anti-LSC activity at 15 nM. Viable CD34+CD38- LSC-enriched **(C)** and CD34- blasts **(D)** relative to DMSO is shown. Active compounds induced LSC depletion and differentiation to CD34- blasts. Mean ± s.d. Representative of *n* = 2 biological replicates. **(E)** Superimposed PDB structures of four corticosteroids from the screen: hydrocortisone (6NWL), triamcinolone acetonide ((5UFS), dexamethasone (1M2Z), and MOM furoate (4P6W), each bound to the glucocorticoid receptor (GR). GR helices are shown in ribbon format, color-coded by root- mean-square deviation. Ligands are shown in aqua/red at the GR binding site. Potent compounds (e.g., MOM) displace GR residue Q111, forming a hydrophobic hotspot that stabilizes receptor binding, whereas inactive compounds (e.g., hydrocortisone) fail to induce this shift. **(F)** Perturbation of the GR ligand-binding pocket by MOM furoate (4P6W). Hydrophobic hotspots are visualized as green patches using Molecular Operating Environment (MOE) electrostatic surface mapping. A 2D interaction map shows the contact points between MOM and the GR hydrophobic surface. **(G)** Summary of predicted binding affinity and conformational changes correlating with anti-LSC activity across screened compounds. HYDRO = Hydrocortisone. TRIAMA = triamcinolone acetonide. DEX = Dexamethasone. MOM = Mometasone.

To investigate how corticosteroid structure influences anti-LSC activity, we screened 24 corticosteroids spanning five structurally defined subclasses (Groups A–D2; Supp. Table S8) in OCI-AML-8227 cells. Fifteen compounds demonstrated anti-LSC activity, with Groups B, C, and D1 exhibiting the highest potency (Figure 4C, Supp. Figure 5A-F). Fluticasone propionate, amcinonide, and triamcinolone acetonide were among the most effective, inducing depletion of the CD34+CD38- LSC-enriched populations and expansion of CD34- blasts at nanomolar concentrations (Figure 4C, D, Supp. Figure 5A-F). Notably, compounds within the same structural subclass displayed similar phenotypic responses, suggesting common structure-activity relationships (Figure 4C, D, Supp. Figure 5A-F). Comparison of active and inactive compounds identified several common features associated with anti-LSC activity, including a C1-C2 double bond, fluorine substitutions at C6/C9, C16 methyl groups, and bulky C17 esters (Supp. Figure 6A- F, Table S9).

To explore the structural basis of these observations, we performed molecular modeling of four representative corticosteroids bound to GR obtained from the Protein Data Bank (PDB) (Figure 4E). Modeling predicted that compounds containing bulky D-ring substituents, such as MOM, displace the Q111 residue from the GR binding pocket, creating hydrophobic hotspots that stabilized ligand binding (Figure 4E, F). In contrast, lower activity compounds such as hydrocortisone failed to induce this conformational change and instead formed weaker hydrogen bonds with Q111 (Figure 4E, F). Importantly, this predicted Q111 conformational shift correlated with anti-LSC activity across the four corticosteroids (Figure 4G). Together, these findings demonstrate that GR engagement is required for GC-induced LSC differentiation in OCI-AML- 8227 cells and identify corticosteroid structural features associated with enhanced anti-LSC potency.

### MOM remodels inflammatory, metabolic and proliferation-associated transcriptional programs in LSCs prior to differentiation

To investigate the early molecular changes that precedes LSC depletion following GC treatment, we performed bulk RNA-sequencing on Fluorescence Activated Cell Sorted (FACS- sorted) CD34+CD38- LSC-enriched OCI-AML-8227 cells, CD34- blasts, and bulk (unsorted) populations following 12 hours of treatment with DMSO control or MOM (0.5 and 10 nM). The 12-hour timepoint was selected to capture early transcriptional responses prior to detectable phenotypic changes (Figure 5A). Unsupervised clustering revealed clear segregation of FACS- sorted CD34+CD38- LSC-enriched cells from CD34- blasts and bulk populations, demonstrating distinct transcriptional responses across cellular compartments (Supp. Figure 7A). Across all six conditions, we identified 988 differentially expressed genes (DEGs) (Figure 5B, Supp. Figure 7B- G, Tables S10–12). Although overlaps were observed, each condition displayed unique DEGs (Supp. Figure 7H-K, Table S10-13).

**Figure 5:**
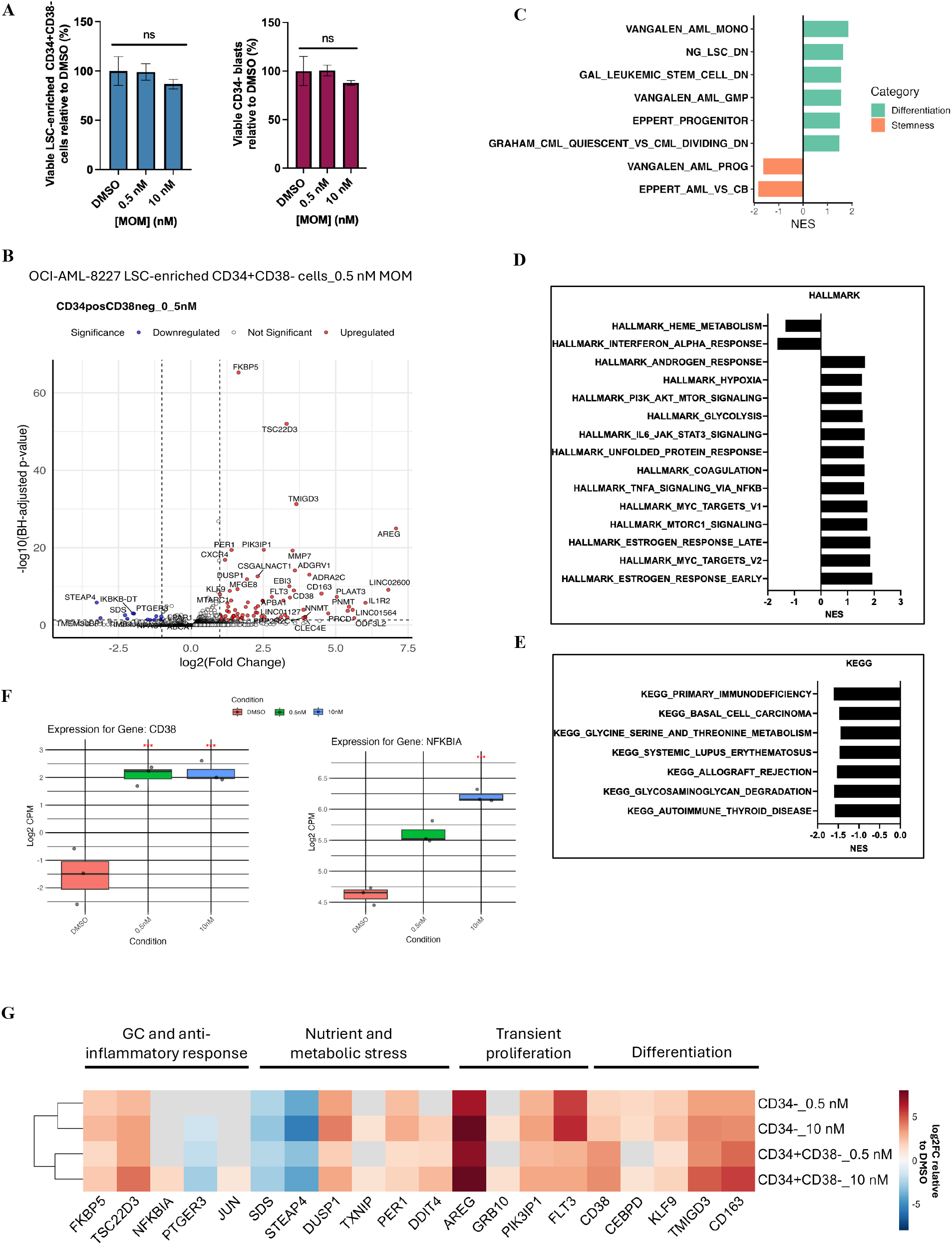
MOM remodels inflammatory, metabolic, and proliferation programs in LSCs prior to differentiation. **(A)** Viable FACS-sorted (left) LSC-enriched CD34+CD38- and (right) CD34- blasts populations 12-hours post-MOM treatment prior to RNA-seq. Demonstrates phenotypic changes are not observed by flow cytometry prior to genomic changes. Mean ± s.d. Representative one of *n* = 3 biological replicates. **(B)** Volcano plot of RNA-seq DEG results from FACS-sorted CD34+CD38- LSC-enriched fraction treated with 0.5 nM MOM for 12 hours vs. DMSO. Genes with |log₂FC| > 1 and BH-adjusted p-value < 0.05 are colored red (upregulated) or blue (downregulated). Labeled genes reflect glucocorticoid response elements, stress, inflammatory, proliferative, and differentiation-related genes. Representative of *n* = 3 biological replicates. **(C)** Targeted GSEA of RNA-seq data from FACS-sorted CD34+CD38- LSC-enriched fractions (12 h post-treatment, 0.5 nM MOM) shows significant negative enrichment of stemness- related signatures and positive enrichment of signatures associated with differentiation, indicating early transcriptional induction of maturation. NES = normalized enrichment score. Significance defined as nominal p < 0.05 and FDR < 0.25. Representative of *n* = 3 biological replicates. **(D)** GSEA of Hallmark pathways in FACS-sorted CD34+CD38- LSC-enriched fraction treated with MOM at 0.5 nM. Pathways have p < 0.05 and FDR < 0.25 significance. Positive enrichment in pathways associated with GC signaling, proliferation and stress in CD34+CD38- LSC-enriched fraction is shown, while inflammatory and nutrient pathways are negatively enriched. Representative of *n* = 3 biological replicates. **(E)** GSEA of KEGG pathways in FACS-sorted CD34+CD38- LSC-enriched fraction treated with MOM at 0.5 nM. Pathways have p < 0.05 and FDR < 0.25 significance. Negative enrichment in pathways associated with inflammation and nutrient metabolism. Representative of *n* = 3 biological replicates. **(F)** Boxplots of certain genes corresponding to the categories in **(D-E)** enriched after 12-hour MOM treatment (0.5 and 10 nM) in FACS-sorted CD34+CD38- LSC-enriched fraction. Log_2_CPM is shown. BH-adjusted p-value was used for significance: * p < 0.05, ** p < 0.01, *** p < 0.001 and **** p < 0.0001. NES: Normalized enrichment score. Representative of *n* = 3 biological replicates. **(G)** Heatmap summarizing key genes from multiple pathways grouped into functional categories, including GC and anti-inflammatory signaling, metabolic stress, transient proliferation, and differentiation. Data show log₂FC relative to DMSO for both FACS-sorted CD34+CD38- LSC-enriched and CD34- blasts fractions. Genes have p < 0.05 and FDR < 0.001 significance. Representative of *n* = 3 biological replicates.

To determine whether these early transcriptional changes were consistent with loss of LSC identity, we performed targeted GSEA on the FACS-sorted CD34+CD38- LSC-enriched fraction. MOM treatment reduced the expression of key LSC maintenance signatures while enriching gene sets associated with progenitor and differentiated myeloid populations (Figure 5C, Supp. Table S14). Several transcriptional changes were additionally consistent with reduced stemness and altered metabolic programs. Expression of *PTGER3*, a gene linked to HSC self-renewal, was downregulated following MOM treatment, while pathways involved in glycine, serine and branched-chain amino acid metabolism were significantly downregulated (Figure 5B,D,E)^38,39^. Additionally, several differentiation-associated genes were upregulated in the FACS-sorted CD34+CD38- LSC-enriched cells, including *CD38*, *CEBPD*, *KLF9*, *TMIGD3*, and *CD163* (Figure 5B, F, G, Supp. Table S12), most of which also increased in the FACS-sorted CD34- blasts (Figure 5G, Supp. Table S11, 16-17). Together, this data demonstrates that the transcriptional commitment toward differentiation precedes the LSC depletion detected by flow cytometry (Figure 5A).

Consistent with a broad remodeling of LSC-associated programs, GSEA of MOM treated FACS-sorted CD34+CD38- LSC-enriched cells revealed significant downregulation of inflammatory and metabolic pathways, including interferon-α response and amino acid metabolism, alongside upregulation of glucocorticoid signaling and stress response pathways such as unfolded protein response and hypoxia (Figure 5D, E, Supp. Table S15). Canonical glucocorticoid target genes, including *FKBP5*, *TSC22D3*, and *NFKBIA,* were among the most highly induced genes in MOM treated FACS-sorted CD34+CD38-LSC-enriched cells (Figure 5B, F, G, Supp. Table S12). Notably, *NFKBIA*, a direct inhibitor of NF-κB signaling, was selectively upregulated in the LSC-enriched fraction and validated by qRT-PCR (Figure 5F-G, Supp. Figure 7L-M), supporting suppression of inflammatory signaling in LSCs following MOM treatment.

MOM treatment also decreased expression of metabolic genes such as *SDS* and *STEAP4* and increased expression of stress-response genes including *DUSP1*, *PER1*, and *DDIT4*, (Figure 5B,G, Supp. Table S12), indicating activation of cellular stress-response pathways.

Interestingly, MOM treatment induced proliferation-associated transcriptional programs in FACS-sorted CD34+CD38- LSC-enriched cells, including enrichment of PI3K, MAPK and RAS signaling pathways and increased expression of proliferation-associated genes *FLT3*, *AREG*, *GRB10*, and *PIK3IP1* (Figure 5B, D, E, G, Supp. Table S12). Similar changes were observed in the FACS-sorted CD34⁻ blasts (Figure 5F, Supp. Table S11, 16), suggesting coordinated transcriptional remodeling or priming across both compartments following GC treatment.

Together, these findings support a model that to reduce stemness, MOM rapidly suppresses inflammatory programs, remodels metabolic pathways and induces proliferation and differentiation associated transcriptional programs in LSCs prior to their phenotypic differentiation and depletion.

### Single cell RNA-sequencing at 24 hours reveals stress-induced differentiation driven by mitochondria hyperactivation and lysosomal dysfunction programs

To determine how the early transcriptional changes observed 12 hours after GC treatment progressed over time, we performed single-cell RNA sequencing (scRNA-seq) of enriched CD34+ LSPCs from OCI-AML-8227 following 24 hours of MOM treatment. Consistent with the flow cytometry data, CD34- blasts emerged by 24 hours post-treatment, resulting in an approximate 60:40 ratio of CD34+ to CD34- cells (Supp. Figure 8A), indicating a shift away from a primitive phenotype.

Using Automated Neural Network Classification for AML Single-Cell Transcriptomes (ANNCAST) and UMAP visualization, we identified seven transcriptionally distinct cell clusters (Figure 6A). MOM treatment resulted in a decrease of the LSC cluster (Figure 6B), consistent with the reduction in the LSC-enriched population observed by flow cytometry (Supp. Figure 8B). Differential expression analysis of the LSC cluster revealed a transcriptional shift consistent with differentiation (Figure 6C, Supp. Table S18). Canonical stemness-associated genes, including *CD34* and *HES1*, were significantly downregulated, whereas hematopoietic regulators including *MPL* and *HEMGN* were strongly upregulated (Figure 6D). Additional differentially expressed genes were associated with oxidative stress, mitochondrial function, lysosomal pathways and inflammation (Figure 6C). Of these, *TXNIP*, *DDIT4*, *NNMT*, *AREG*, and *FKBP5* were also upregulated at 12 hours, indicating persistence of the early glucocorticoid response (Figure 5F). In contrast, most transcriptional changes were unique to the 24-hour timepoint, suggesting progression from the early stress response toward a differentiation-associated state.

**Figure 6:**
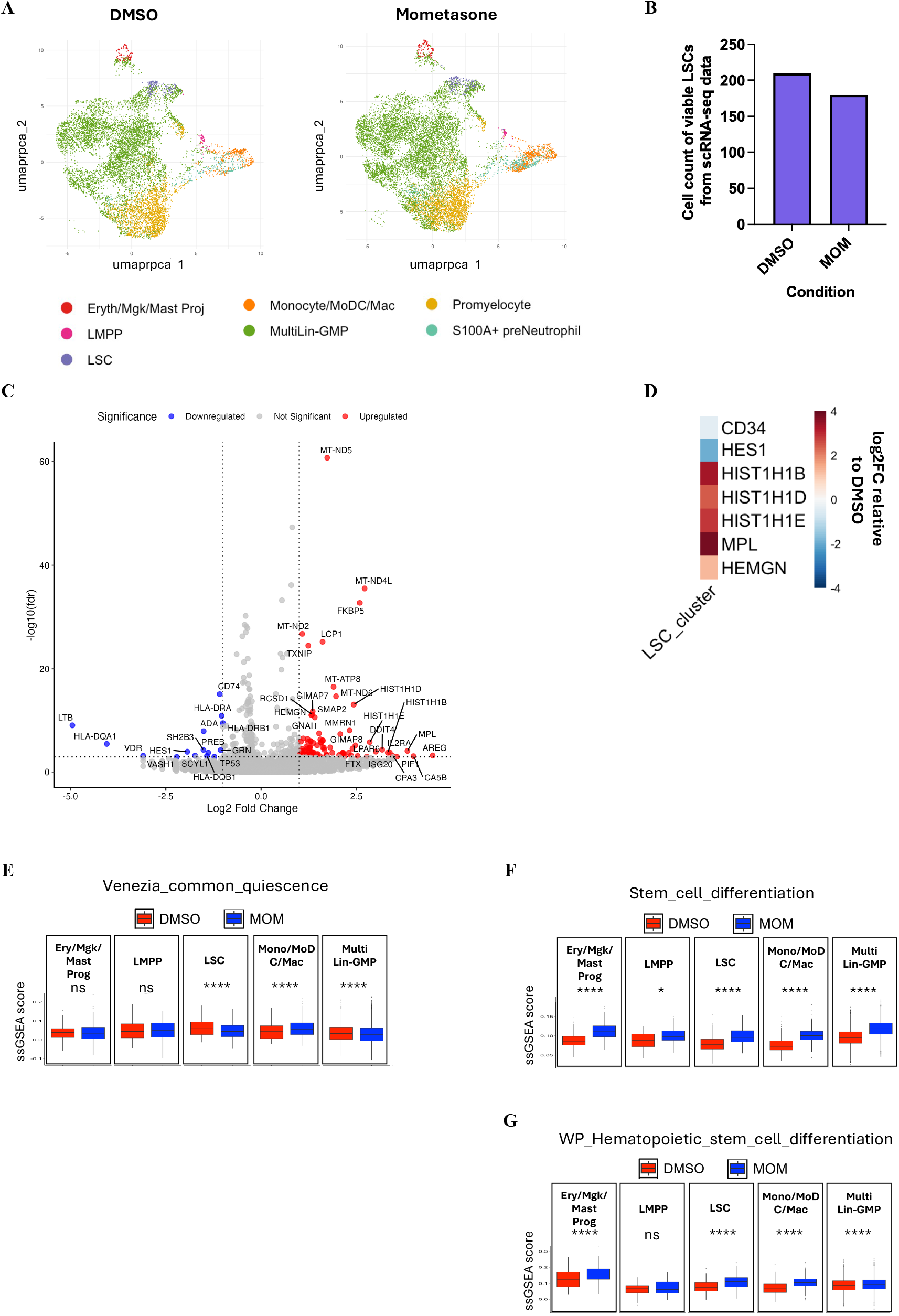
Single-cell RNA-seq reveals MOM-induced loss of leukemic stemness and promotion of differentiation programs at 24 hours. **(A)** UMAP visualization of scRNA-seq data from enriched OCI-AML-8227 CD34+ LSPCs at 24 hours post-treatment with DMSO (left) or 0.5 nM of MOM (right). Clusters were assigned using the ANNCAST v1.2 classifier, identifying seven transcriptional populations corresponding to HSPCs and their downstream progeny. The LSC cluster (purple) is markedly depleted upon treatment, with a corresponding increase in leukemic progenitor populations and differentiated CD34- blasts, including MultiLin GMP (green), Monocyte/MoDC/Mac (orange), and Promyelocytes (yellow). *N* = 1 biological replicate for scRNA-seq data. **(B)** Quantification of viable LSC cell count from LSC cluster scRNA-seq data is shown for each condition. **(C)** Volcano plot showing differentially expressed genes (DEGs) in the LSC cluster (CD34+CD38-) following 24-hour MOM treatment normalized to DMSO. A cutoff of |log₂ fold change| > 1 and FDR < 0.001 was used to define significance. Genes meeting this threshold and downregulated are colored in blue, upregulated genes in red, and non-significant genes in grey. **(D)** Heatmap showing expression of key genes associated with stemness (*CD34*, *HES1*) and differentiation (*HIST1H1B*, *HIST1H1D*, *HIST1H1E*, *MPL*, *HEMGN*) in the LSC cluster at 24 hours. **(E-G)** ssGSEA analysis of all seven cluster types indicate **(E)** negative enrichment of HSPC quiescence-related signature and **(F, G)** positive enrichment of stem cell differentiation signatures in LSC cluster. Wilcoxon test comparing the scores between DMSO and MOM for each cell type was performed: * < 0.05, ** < 0.01, *** < 0.001 and **** < 0.0001. Representative of *n* = 1 biological replicate.

To quantify changes in cell state, we performed single-sample GSEA (ssGSEA). For the LSC cluster, significant depletion of HSC quiescence signatures together with enrichment of hematopoietic stem cell, progenitor and myeloid differentiation programs were observed (Figure 6E-G, Supp. Figure 8C-F, Table S19). Compared with the 12-hour bulk RNA-seq data, proliferation-related programs were less prominent, with only HALLMARK_MITOTIC_SPINDLE remaining enriched, suggesting that proliferation-associated transcriptional programs are transient following MOM treatment (Figure 7A).

**Figure 7:**
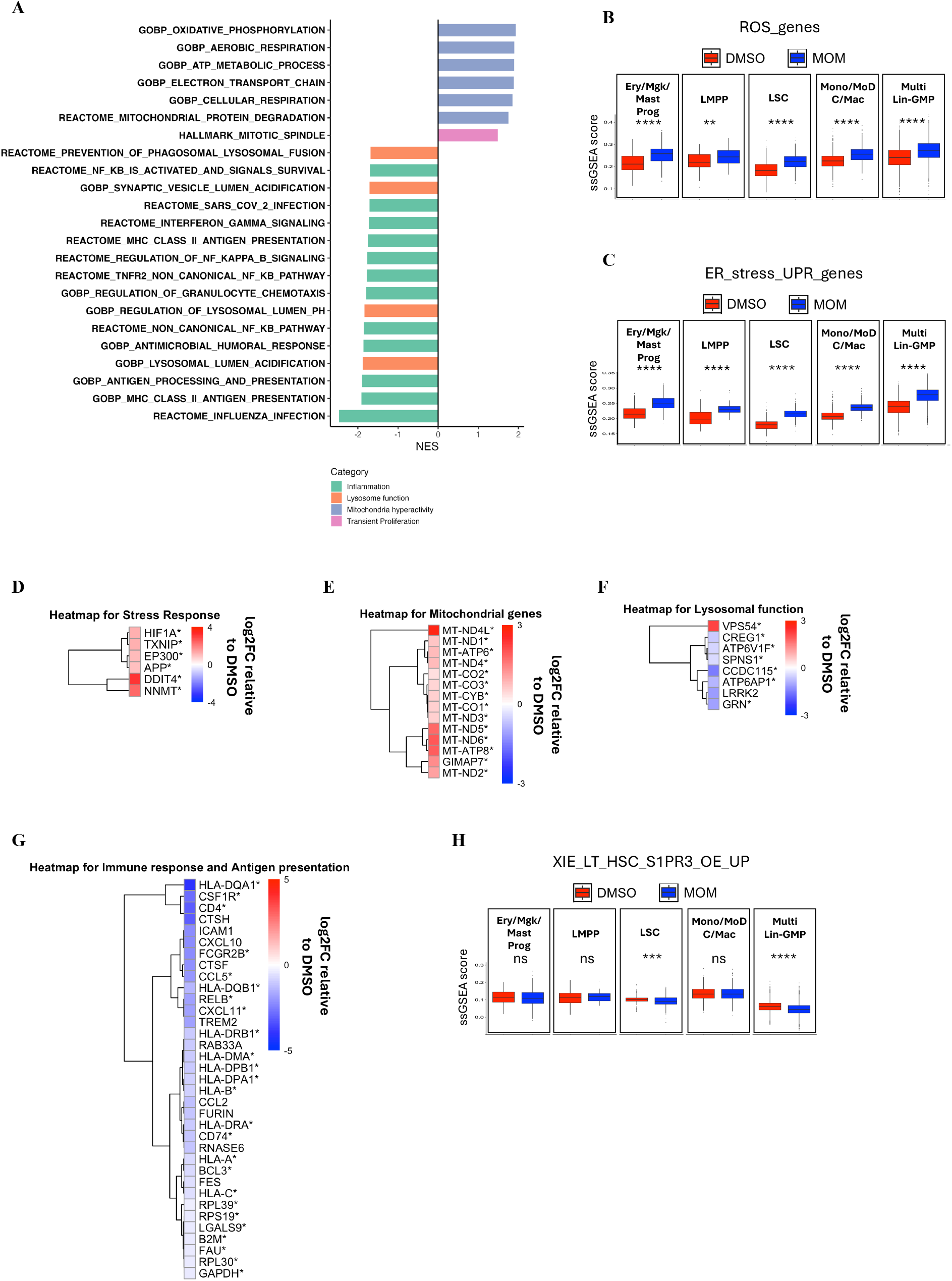
Single-cell RNA-seq at 24 hours reveals GC stress-induced differentiation in LSCs is driven by mitochondria hyperactivation, lysosomal dysfunction and continued suppression of immune programs. **(A)** GSEA of the LSC cluster (MOM vs. DMSO, 24 h) using curated signatures from the GO Biological Process (GOBP) and Reactome databases. Pathways with nominal p-value < 0.05 and FDR < 0.25 are shown. NES = normalized enrichment score. Enriched gene sets reflect persistent oxidative and mitochondrial stress (blue), lysosomal dysfunction (orange), and suppression of inflammatory and antigen presentation pathways (green). The mitotic spindle program (pink) is the only remaining proliferation signature, suggesting exit from a transient proliferative state. **(B-C)** ssGSEA demonstrates **(B)** ROS_genes and **(C)** ER_Stress_UPR_genes signatures are positively enriched in the LSC cluster. These signatures indicate sustained oxidative stress, unfolded protein and ER stress. Wilcoxon test comparing the scores between DMSO and MOM for each cell type was performed: * < 0.05, ** < 0.01, *** < 0.001 and **** < 0.0001. **(D-G)** Heatmaps of leading-edge genes from significant GSEA signatures. Log-normalized gene expression in the LSC cluster is shown (MOM vs. DMSO, 24 h). Genes with an asterisk are significant in DEG data and have p < 0.05. **(D)** Stress response genes are upregulated, confirming activation of oxidative and metabolic stress programs. (**E)** Electron transport chain genes are significantly upregulated, indicating mitochondrial hyperactivation. **(F)** Lysosomal and autophagy-related genes are mainly downregulated, consistent with lysosomal dysfunction and impaired autophagic flux. **(G)** Immune regulatory and MHC class II antigen presentation genes are broadly suppressed, reflecting sustained repression of inflammatory signaling. **(H)** ssGSEA of the LT_HSC_S1PR3_OE_UP signature from Xie *et a*l., 2021. This inflammation-driven long-term HSC signature is negatively enriched in the LSC cluster after treatment, supporting a model of glucocorticoid-induced differentiation that is mechanistically distinct from TNF-a-driven inflammatory differentiation. Wilcoxon test: *<0.05, **<0.01, *** < 0.001, **** < 0.0001. *N* = 1 biological replicate for scRNA-seq data and analysis.

Stress-associated pathways demonstrated persistent enrichment of reactive oxygen species (ROS), endoplasmic reticulum stress and unfolded protein response programs at 24 hours in the LSC cluster (Figure 7A-C Supp. Table S19). This was accompanied by increased expression of multiple stress response genes (Figure 7D). Mitochondrial pathways were also positively enriched (Figure 7A, Supp. Table S19), with upregulation of genes associated with electron transport chain components (Figure 7A, E, Supp. Table S19). In parallel, lysosomal and autophagy-associated pathways were negatively enriched (Figure 7A, Supp. Table S19), with consistent downregulation of multiple pathway-associated genes (Figure 7F). Together, these findings indicate persistent remodeling of metabolic and stress transcriptional programs following GC treatment.

Robust suppression of immune and inflammatory pathways remained at 24 hours, extending the anti-inflammatory response observed from bulk RNA-seq at 12-hours post-treatment (Figure 5E and 7A). Downregulation of NF-κB signaling, MHC class II pathways and additional immune related programs were observed in the LSC cluster (Figure 7G, Supp. Table S19). This mechanism is distinct from TNF-α–induced inflammation-driven differentiation, as shown by the negative enrichment of the Xie *et al*. 2021 long-term HSC signature and upregulation of *S1PR3* (Figure 7H)^40^ .Together, these findings define a mechanistic model in which stress-induced metabolic dysfunction and consistent repression of inflammatory signaling converge to drive LSC differentiation and CD34- blast expansion.

### FLT3 ligand supports CD34- blast expansion but is dispensable for LSC depletion

Bulk RNA-seq analyses revealed induction of proliferation transcriptional programs following MOM treatment at 12 hours, including increased expression of *FLT3* in the LSC- enriched population at 12 and 24 hours (Figure 5 and 8A, Supp. Table S12, 18). We therefore investigated whether specific cytokines are required for the proliferative response associated with GC-induced differentiation. OCI-AML-8227 cells were cultured in the absence of individual growth factors (SCF, TPO, IL-3, FLT3L, IL-6 or G-CSF) and treated with increasing concentrations of MOM (Figure 8B). Depletion of the LSC-enriched population occurred across all conditions, demonstrating that GC-mediated LSC depletion does not require any individual growth factor (Figure 8C). In contrast, expansion of CD34- blasts was selectively dependent on FLT3L and was significantly decreased under FLT3L deficient conditions (viable CD34- blasts relative to DMSO with all growth factors vs. –FLT3L at 1000 nM MOM: 203% vs. 96.5%) (Figure 8C).

**Figure 8:**
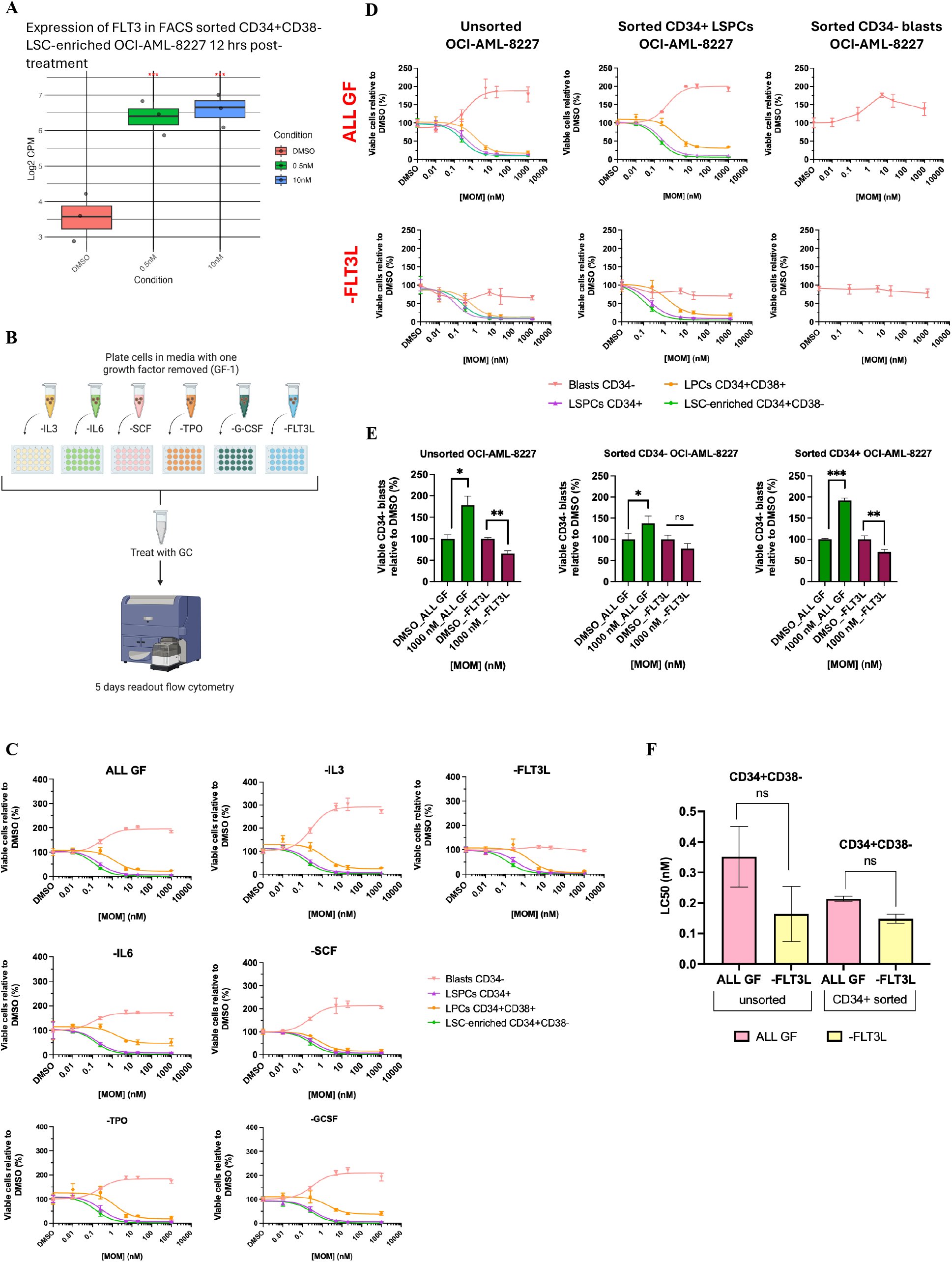
FLT3L is required for CD34- blast expansion but not for LSC-directed differentiation. **(A)** Boxplot of FLT3 log_2_CPM gene expression change in FACS-sorted CD34+CD38- LSC-enriched OCI-AML-8227 cells 12 hours post-MOM treatment (0.5 and 10 nM). Data from bulk RNA-seq. BH-adjusted p-value was used for significance: * p < 0.05, ** p < 0.01, *** p < 0.001 and **** p < 0.0001. Representative of *n* = 3 biological replicates. **(B)** Schematic of the growth factor minus 1 (GF-1) experiment: OCI-AML-8227 cells were plated in media with one cytokine removed per condition (IL-3, IL-6, SCF, TPO, G-CSF, or FLT3L), treated with increasing doses of MOM and analyzed by flow cytometry after five days. **(C)** Dose-response curves for each growth factor–depleted condition. Viable cells relative to DMSO control. Mean ± s.d. Representative one of *n* = 5 biological replicates. **(D)** Unsorted cells, FACS-sorted CD34+ LSPCs and CD34- blasts from OCI-AML-8227 were treated with increasing doses of MOM in the presence of all growth factors (ALL GF) or in the absence of FLT3L for five days. Viable cells relative to normalized to DMSO control is shown. Mean ± s.d. Representative one of *n* = 3 biological replicates. **(E)** Viable CD34- blasts counts relative to DMSO in unsorted, CD34- FACS- sorted blasts, and FACS-sorted CD34+ LSPCs from OCI-AML-8227 cells cultured in the presence or absence of FLT3L and treated with DMSO or 1000 nM MOM. *p < 0.05, **p < 0.01, **p < 0.001; unpaired t-test; mean ± s.d. Representative one of *n* = 3 biological replicates. **(F)** LC₅₀ values for CD34+CD38- LSC-enriched fraction treated with MOM from unsorted or FACS-sorted CD34+ LSPCs cultured in the presence or absence of FLT3L. Mean ± s.d. Representative one of *n* = 4 biological replicates. ns = non-significant.

We next asked whether the FLT3 dependent expansion of CD34- cells resulted exclusively from differentiation of CD34+ LSPCs or whether pre-existing CD34- blasts could also expand in response to MOM treatment. To test this, we FACS-sorted CD34+ and CD34- fractions from OCI- AML-8227 and repeated the assay in the presence or absence of FLT3L. Following MOM treatment, purified CD34+ LSPCs generated CD34- blasts, consistent with differentiation from the LSC-enriched populations (Figure 8D). Expansion of these newly generated CD34- blasts was observed only in the presence of FLT3L but not in the absence (viable CD34- blasts relative to DMSO with ALL GF vs. –FLT3L at 1000 nM MOM: 192% vs. 70%, p = 0.007) (Figure 8D). Similarly, FACS-sorted CD34- cells expanded following MOM treatment only when FLT3L was present (viable CD34- blasts relative to DMSO at 1000 nM MOM for ALL GF: 138%, p = 0.04, and for -FLT3L: 78%, p = 0.06, respectively) (Figure 8D, E). Together, these findings indicate that FLT3L selectively supports expansion of the CD34- blast population following GC treatment.

To determine whether FLT3L influences GC sensitivity in LSCs, we calculated LC₅₀ values for the CD34+CD38- LSC-enriched population in the presence or absence of FLT3L in unsorted and FACS-sorted conditions (Figure 8F). No significant differences in LC₅₀ were observed across conditions (Figure 8F), confirming that FLT3L is dispensable for MOM depletion of the LSC-enriched population.

Together, these findings demonstrate that FLT3L is required for the blast expansion associated with GC treatment while being unnecessary for the depletion of the LSC-enriched population.

### Glucocorticoid treatment depletes functional LSCs and reprograms AML cells toward a differentiated state

To determine whether the phenotypic loss and transcriptional suppression of stemness- associated programs following *in vitro* treatment by MOM translates into functional loss of leukemia stem cell activity, we performed xenotransplantation assays using OCI-AML-8227 cells treated with MOM or DMSO for seven days followed by intrafemorally injections into NSG-S mice (Figure 9A, B). After 12 weeks, human AML engraftment in the injected femur was significantly reduced in the MOM-treated group (p < 0.0001), as well as in the contralateral femur (p < 0.001) and spleen (p < 0.01) (Figure 9C). These findings demonstrate that MOM treatment substantially reduces the leukemia-initiating capacity of LSCs in human AML.

**Figure 9:**
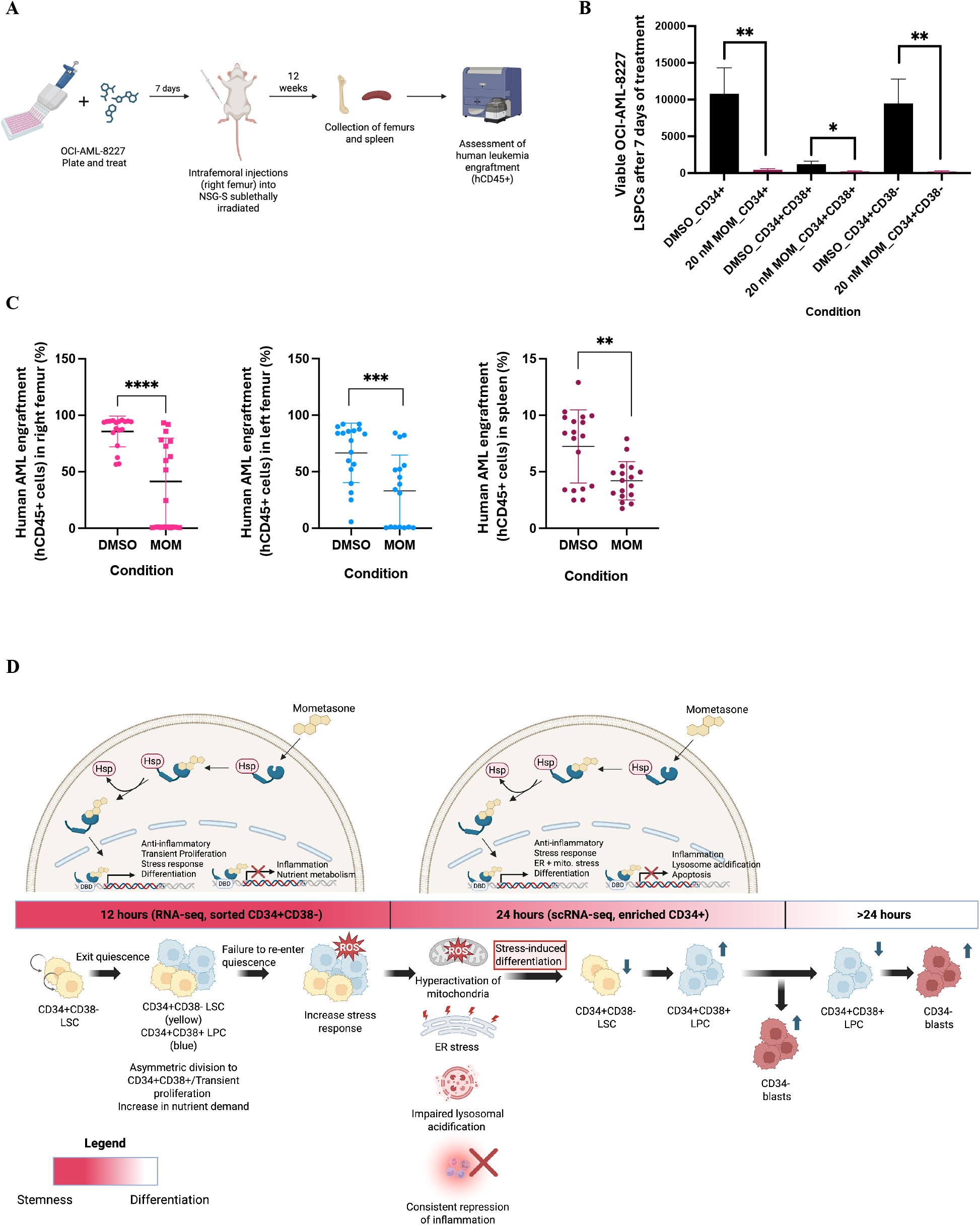
MOM-induced differentiation of leukemic stem cells reduces leukemia-initiating capacity *in vivo*. **(A)** Experimental schematic for *in vivo* leukemia-initiating potential assay. Bulk OCI-AML-8227 cells were treated with 20 nM MOM for seven days and 1/4 of total cells were injected intrafemorally into sublethally irradiated NSG-S mice (2.1 Gy). After 12 weeks, femurs and spleens were harvested for analysis by flow cytometry. **(B)** Viable CD34+ LSPCs and CD34- blast cell counts following seven-day *in vitro* treatment with 20 nM MOM, prior to injection in NSG-S mice. Mean ± s.d. unpaired t-test: *p<0.05, **p<0.01, ***p<0.001, ****p<0.0001. Representative one of *n* = 3 biological replicates. **(C)** Human AML leukemia engraftment (hCD45+) in injected femur (left), contralateral femur (middle), and spleen (right) at 12 weeks. Each point represents an individual mouse. Representative of *n* = 3 biological replicates, *n* = 6 mice per condition (one mouse was removed from one biological replicate for MOM condition as it did not meet the endpoint). Mean ± s.d. is shown. Mann-Whitney U test was used for statistical testing: *p<0.05, **p<0.01, ***p< 0.001, ****p< 0.0001. **(D)** Working model of glucocorticoid-driven LSC differentiation. At 12 hours, CD34+CD38- LSCs exit quiescence due to suppressed inflammation and elevated stress signaling. By 24 hrs, mitochondrial hyperactivation, ER stress, and lysosomal dysfunction trigger stress-induced differentiation, while inflammatory signaling remains repressed. This cascade results in the progressive depletion of LSCs and LPCs, with an increase in CD34- blast output and loss of leukemia-initiating potential.

Together with the phenotypic and transcriptional data, these findings demonstrate that MOM treatment depletes CD34+CD38- LSCs through reprogramming inflammatory, metabolic and lysosomal programs to suppress stemness-associated programs and promote differentiation to CD34- blasts (Figure 9D). Collectively, these results support the development of glucocorticoids as differentiation inducing therapies targeting the LSC compartment in AML.

## DISCUSSION

AML remains a clinical challenge due to the persistence of LSCs, which evade conventional therapies and drive relapse. In this study, we demonstrate that GCs induce rapid depletion of AML LSCs by inducing terminal differentiation. Unlike their pro-apoptotic role in lymphoid malignancies, GCs promoted loss of stemness, triggered differentiation and reduced leukemia-initiating capacity in AML. Importantly, these effects were observed across multiple LSC-containing models and genetically diverse primary AML samples.

Mechanistically, GC-induced LSC depletion through differentiation was dependent on glucocorticoid receptor signaling. Pharmacologic antagonism using RU486 blocked both differentiation and LSC depletion, demonstrating a requirement for GR activation. Furthermore, corticosteroid screening and structure-activity modeling identified specific structural features associated with enhanced anti-LSC activity, including the C1-C2 double bond and bulky D-ring substituents. These findings are consistent with previous studies in non-hematopoietic systems that showed such substitutions enhance GR binding and reduce off-target effects through stabilization of the hydrophobic ligand-binding pocket^42^. To our knowledge, our work is the first to demonstrate that these structural features correlate with anti-LSC activity in primary human AML models, identifying MOM as a particularly potent candidate for therapeutic development.

A major finding of this study is that GC responsiveness was not associated with recurrent AML mutations or baseline *NR3C1* expression. Although previous reports emphasized specific genetic contexts, including RUNX1- and NPM1-mutated AMLs^24,26^, our data indicates that no single mutation consistently dictated glucocorticoid responsiveness, and transcriptional state prior to treatment is a stronger predictor than genotype. AML samples with enriched inflammatory programs preferentially underwent differentiation whereas samples enriched in extracellular matrix programs more commonly responded through apoptosis. Consistent with this observation, OCI-AML-8227, which undergoes GC-induced differentiation, exhibited high baseline inflammatory gene expression, suggesting that inflammation-prone LSCs are particularly vulnerable to GC-mediated immune suppression. These findings support a model in which cellular state, rather than mutational profile alone, may govern therapeutic susceptibility. Although 21 samples were investigated, larger cohorts would need to be used to validate this finding in the future.

Among the transcriptional changes induced by GC treatment, suppression of inflammatory signaling emerged as a central feature. Baseline inflammatory programs were enriched in differentiation prone AML samples, while GC treatment induced *NFKBIA* and other regulators that repress NF-κB signaling. Given the established role of NF-κB in maintaining AML stemness and survival^18^, these findings suggest that GCs target a key dependency of LSCs. Unlike prior attempts to target this pathway pharmacologically (e.g., with parthenolide), which has been limited by toxicity and bioavailability^43–45^, GCs represent a clinically established approach capable of suppressing inflammatory programs while promoting differentiation. Furthermore, targeting inflammatory programs to sensitize AML samples to GCs has been supported by previous studies in therapy-resistant AML. Acquisition of resistance to FLT3 inhibitors in FLT3-ITD AMLs or cytarabine in FLT3-wt AMLs has been associated with increased inflammatory signaling and enhanced sensitivities to GCs^29,30^. Similarly, a third study showed that *NPM1*-mutant AMLs resistant to Ara-C had upregulated inflammatory programs and responded to GC-based therapy, reinforcing the observation that inflammation can sensitize AML cells to GCs^26^. Together with our findings, inflammatory transcriptional programs may represent a common determinant of GC responsiveness and could help identify patients most likely to benefit from this therapy.

In addition to inflammatory remodeling, bulk and scRNA-seq analyses of the LSC- enriched population identified activation of stress-response pathways together with extensive remodeling of metabolic programs. We observed enrichment of mitochondrial respiratory pathways and suppression of lysosomal and autophagy-associated programs following GC treatment. Although these observations were derived from transcriptomic analyses and require further functional validation, they are consistent with the emergence of persistent cellular stress accompanying differentiation. LSCs rely on OXPHOS over glycolysis, making them resistant to cytotoxic therapies but vulnerable to metabolic perturbation^12^. Lysosomes play a critical role in maintaining quiescence and regulating stress responses in HSCs^46,47^, and AML LSCs exhibit enlarged lysosomes that have been proposed as a therapeutic vulnerability, including through the inhibition of the V-ATPase complex^48–50^. Thus, GC-mediated lysosomal dysfunction in combination with repression of inflammation may tip LSCs beyond their adaptive capacity, enforcing a differentiation fate. Similar stress-associated differentiation mechanisms have been proposed in other hematologic malignancies, such as in chronic myeloid leukemia LSCs, where inhibition of autophagy induced mitochondrial stress and promoted differentiation^51^. A similar stress-induced model may apply in AML, where GCs may exploit vulnerabilities associated with the metabolic plasticity of AML LSCs but warrants further investigation.

A key translational insight in this study is that while FLT3L is dispensable for the therapeutic effect on LSCs, it may influence the degree of blast proliferation following GC treatment. Since FLT3L levels can rise under inflammatory conditions or chemotherapy, it may be beneficial to monitor FLT3L to predict blast expansion in GC-treated patients and guide clinical management to avoid complications such as leukostasis. In AML, the FLT3 receptor is retained in both LSCs and leukemic blasts, where FLT3L promotes their proliferation and survival^52,53^. Previous studies have demonstrated that FLT3L promotes AML blast proliferation and can enhance GR activity *in vitro* when combined with dexamethasone^54,55^. These findings raise the possibility that FLT3L may cooperate with GCs to drive blast proliferation, warranting clinical caution in high-FLT3L contexts.

Importantly, we show that MOM synergizes with cytotoxic agents like Ara-C, supporting combination strategies that integrate differentiation-inducing and cytotoxic mechanisms. More broadly, our findings add to growing evidence that forcing LSCs out of a stem-like state represents an effective therapeutic strategy in AML. Similar principles were demonstrated with the success of differentiation therapy in APL and emerging approaches targeting IDH1/2, FLT3, BRD4 and menin^56–60^. GCs can join this class, promoting a non-cytotoxic, stress-induced exit from stemness with a strong therapeutic index for non-APL AMLs.

In conclusion, we demonstrate that GCs induce differentiation depletion of functional AML LSCs through NR3C1-dependent transcriptional reprogramming. GC treatment is accompanied by suppression of inflammatory programs, remodeling of metabolic pathways, and acquisition of differentiation cell states, resulting in the reduction of leukemia-initiating capacity. These findings support the clinical development of GCs as differentiation-based therapies targeting AML LSCs and provide a framework for biomarker guided patient stratification in AML.

## Supporting information

Supplementary Table Titles and Legends

## ACKNOWLEDGEMENTS

This work was supported by grants from the Cole Foundation (KE), the Ellie White Fund (KE), CIHR grants (KE), Leukemia and Lymphoma Society (KE) and fellowships from CIHR and FRQS (II). We would like to thank Dr. Guy Sauvageau for his kind donation of the corticosteroids used in this project.

