## Supplementary Figures for "Glucocorticoids reprogram human AML leukemic stem cells to promote elimination through differentiation and apoptosis"

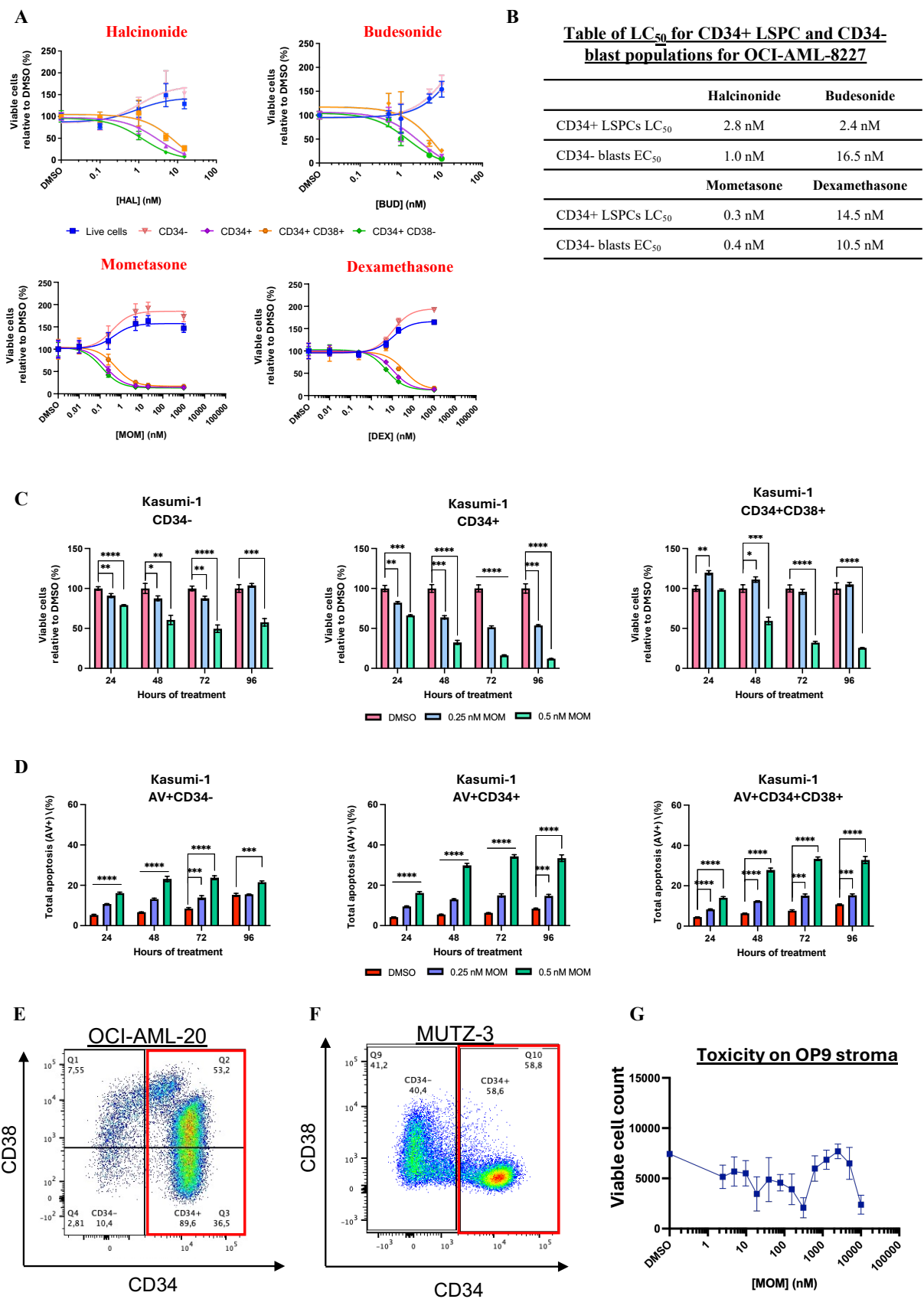

**Supplemental Figure 1: Glucocorticoids induce differentiation without apoptosis in LSC models as early as 24 hours. (A)** Dose response treatment of OCI-AML-8227 with FDA-approved compounds halcinonide, budesonide, MOM and dexamethasone. Differentiation of LSC-enriched fraction CD34<sup>+</sup>CD38<sup>-</sup> and leukemic progenitors CD34<sup>+</sup>CD38<sup>+</sup> (LPCs) to leukemic blasts CD34<sup>-</sup> is observed. Mean ± s.d. Representative one of $n = 3$ biological replicates. **(B)** The LC<sub>50</sub> values for the depletion of CD34<sup>+</sup> LSPC fraction and EC<sub>50</sub> for expansion of CD34<sup>-</sup> blast cells by glucocorticoids. Nonlinear regression fit model using Prism v9 was used with 95% CI. Representative one of $n = 3$ biological replicates. **(C-D)** Kasumi-1 time course treated with MOM (0.25 and 0.5 nM) for 24 to 96 hrs. Cell viability relative to DMSO **(C)** and apoptosis **(D)** are shown for CD34<sup>-</sup>, CD34<sup>+</sup> and CD34<sup>+</sup>CD38<sup>+</sup>. Apoptosis was determined using Annexin V and 7-AAD. P value calculated using unpaired student t-test. Mean ± s.d. Representative one of $n = 3$ biological replicates. **(E-F)** Flow cytometry plots of CD38 vs CD34 LSC-models **(E)** OCI-AML-20 and **(F)** MUTZ-3. Red box highlights CD34<sup>+</sup> LSC-enriched fractions in both models. OCI-AML-20 is a co-culture model requiring OP9 stroma. **(G)** Viable OP9 cell count after five-day treatment with increasing doses of MOM. Mean ± s.d. Representative one of $n = 3$ biological replicates.

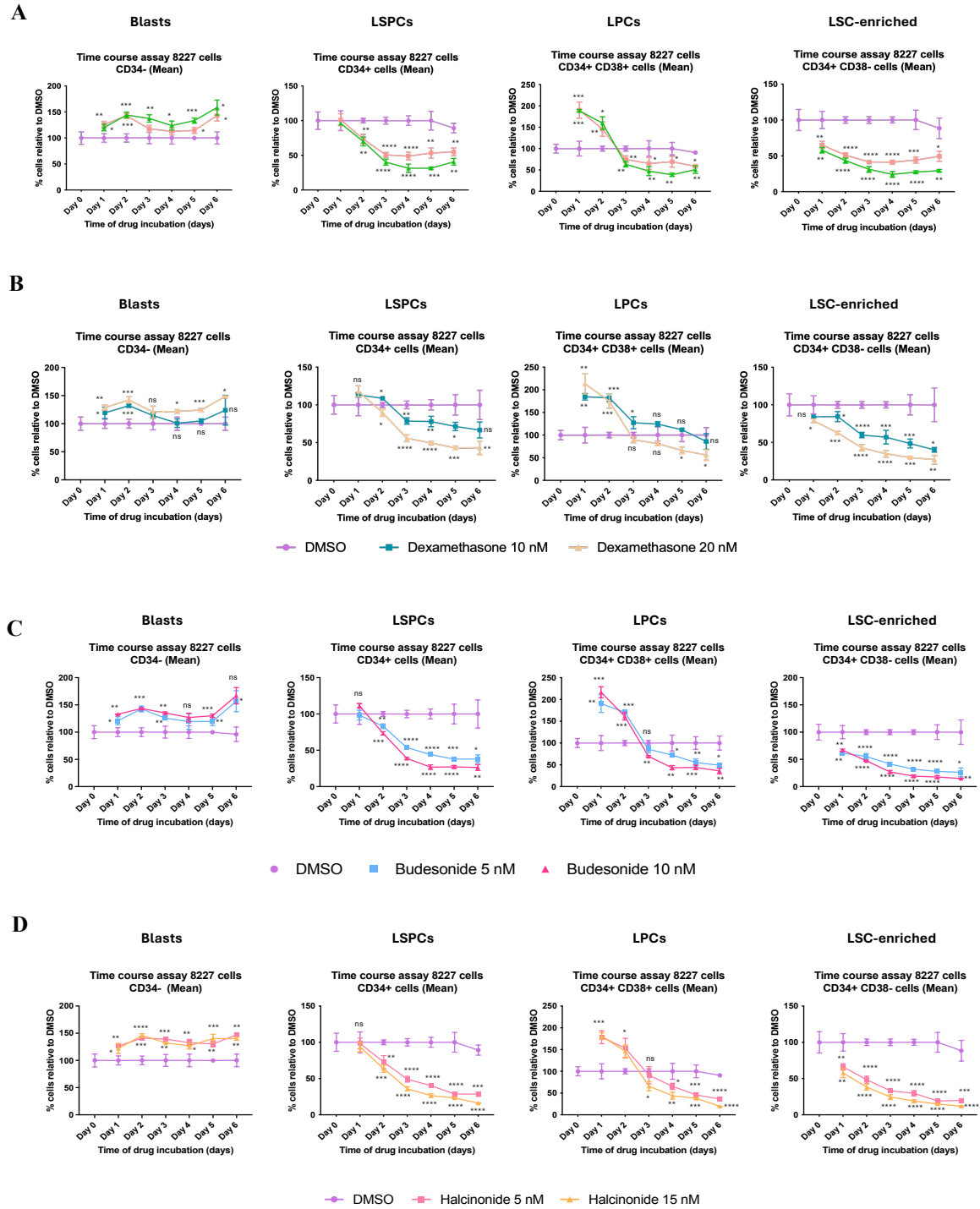

**Supplemental Figure 2: Kinetics of GC-induced differentiation of LSPCs in OCI-AML-8227.**

**(A-D)** Six-day time course assay of OCI-AML-8227 cells treated with all four glucocorticoids **(A)**

**MOM** (0.64 and 1.6 nM), **(B)** dexamethasone (10 and 20 nM), **(C)** budesonide (5 and 10 nM) and

**(D)** halcinonide (5 and 15 nM). Purple line indicates DMSO/untreated control. Cell viability relative to DMSO (indicated as % cells relative to DMSO). Analysis shown for different populations: CD34<sup>-</sup> blasts, CD34<sup>+</sup> LSPC fraction, CD34<sup>+</sup>CD38<sup>+</sup> LPCs and CD34<sup>+</sup>CD38<sup>-</sup> LSC-enriched fraction. One-way ANOVA, \* $p < 0.05$, \*\* $p < 0.01$, \*\*\* $p < 0.001$ and \*\*\*\* $p < 0.0001$. Mean ± s.d. Representative one of $n = 2$ biological replicates.

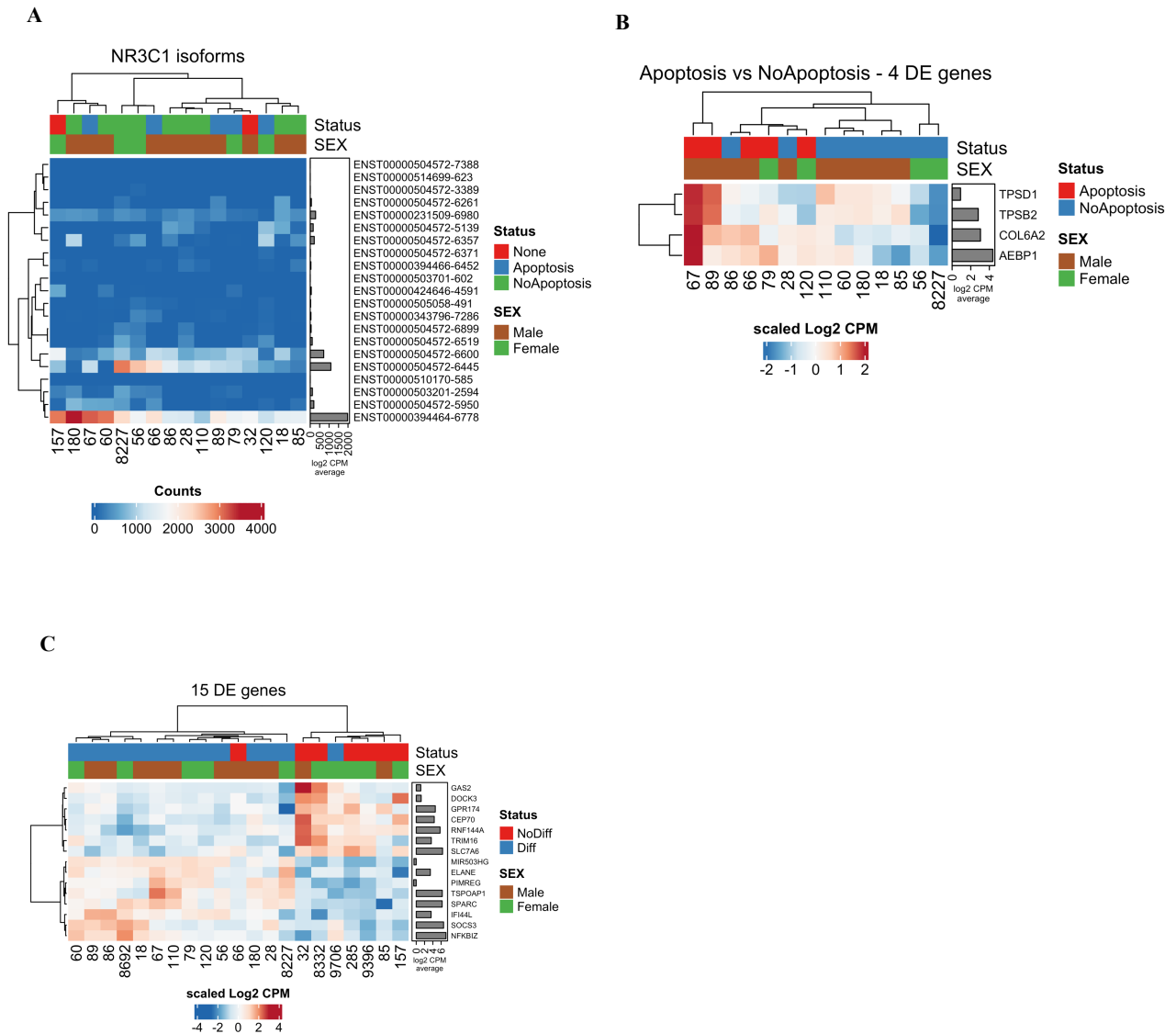

**Supplemental Figure 3: Gene expression programs, but not GR levels or mutations, predict glucocorticoid response in AML.** (A) Heatmap showing correlation between NR3C1 isoforms expression and glucocorticoid response in AML samples. A correlation between GR isoforms and the type of response induced (apoptosis or differentiation) is not observed and therefore not predictive of response. Representative of $n = 1$ biological replicate, $n = 16$ patient samples including OCI-AML-8227. (B) Heatmap of differentially expressed genes in pre-treated AML

samples that responded with apoptosis to MOM. Unsupervised clustering is performed for apoptosis vs no apoptosis response and sex status. Genes are involved in ECM remodeling and immune response, indicating a correlation between ECM signatures and apoptosis. Representative of $n = 1$ biological replicate, $n = 14$ patient samples including OCI-AML-8227. **(C)** Heatmap of differentially expressed genes in pre-treated AML samples that responded with differentiation to MOM vs those that did not. Unsupervised clustering is performed for differentiation vs no differentiation response and sex status. Genes associated with immune cell differentiation, scavenger pathways and inflammation-related pathways are apparent. Representative of $n = 1$ biological replicate, $n = 21$ patient samples including OCI-AML-8227. Red is no response, green is response through differentiation, and no apoptosis and blue is response through apoptosis.

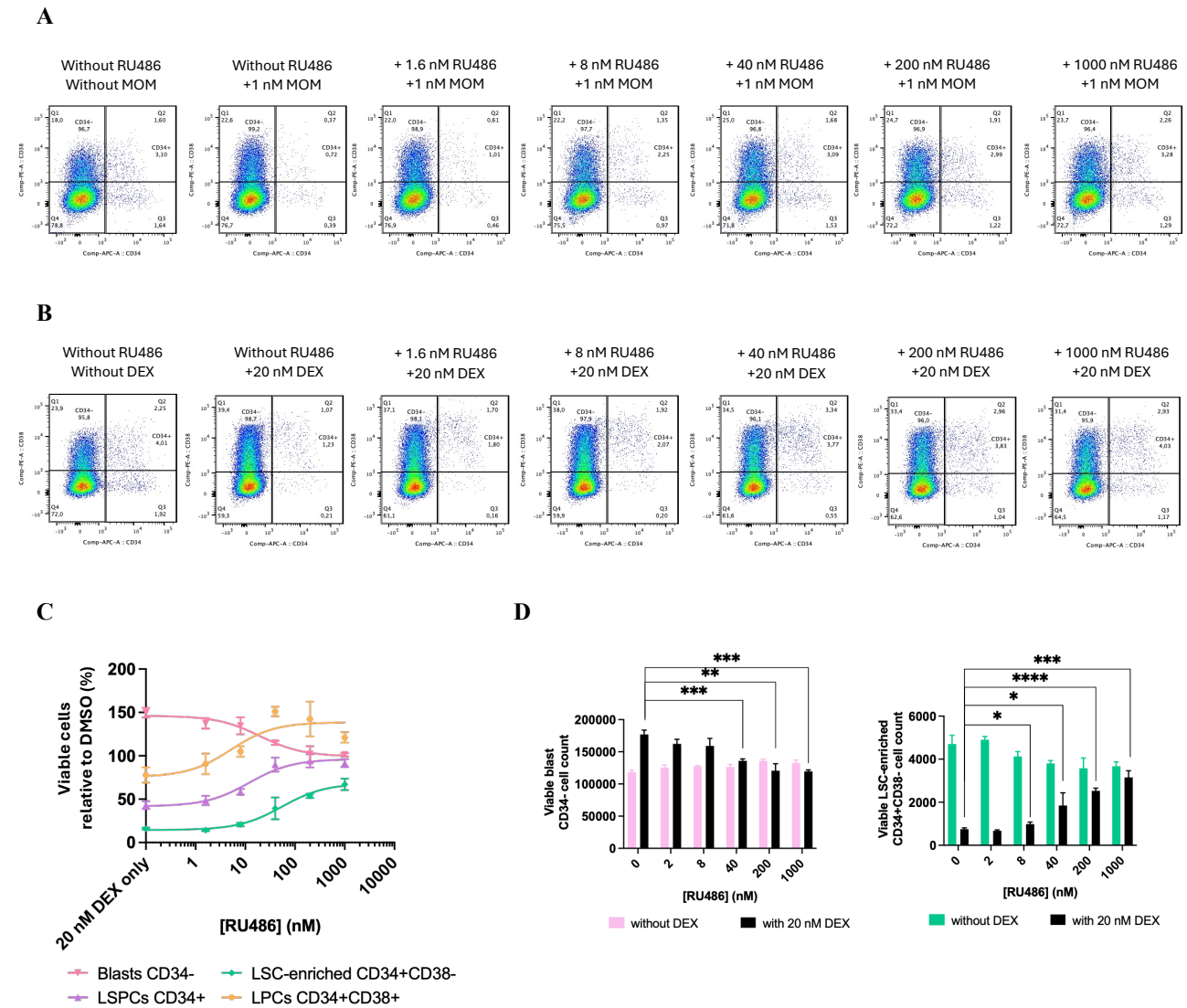

**Supplemental Figure 4: RU486 blocks glucocorticoid mediated LSC differentiation. (A-B)**

Flow cytometry profile of CD38 vs CD34 OCI-AML-8227 cells treated with 1 nM of MOM (A) or 20 nM of dexamethasone (DEX) (B) with increasing concentration of the GR antagonist, RU486. Representative one of $n = 3$ biological replicates. (C) RU486 blocks DEX-induced LSC differentiation. OCI-AML-8227 cells were treated with 20 nM DEX with or without the glucocorticoid receptor (NR3C1) antagonist RU486 for six days. RU486 prevents depletion of the CD34+CD38- LSC-enriched population and CD34- blast expansion, confirming that NR3C1 signaling is required for GC-induced differentiation. Viable cells relative to DMSO is shown.

Mean ± s.d. Representative one of $n = 3$ biological replicates. **(D)** Viable cell counts for CD34-blasts (left) and CD34<sup>+</sup>CD38<sup>-</sup> LSC-enriched fraction (right) for OCI-AML-8227 treated with 20 nM DEX ± RU486. Starting dose of 0 nM for the colored bar indicates absence of RU486, for black bar indicates 20 nM of DEX is present only. RU486 is then added at increasing doses. Unpaired t-test, \* $p < 0.05$, \*\* $p < 0.01$, \*\*\* $p < 0.001$ and \*\*\*\* $p < 0.0001$. Mean ± s.d. Representative one of $n = 3$ biological replicates.

A

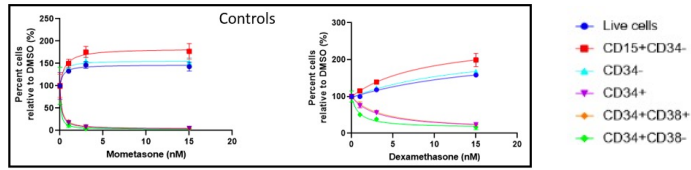

B

**Group A**

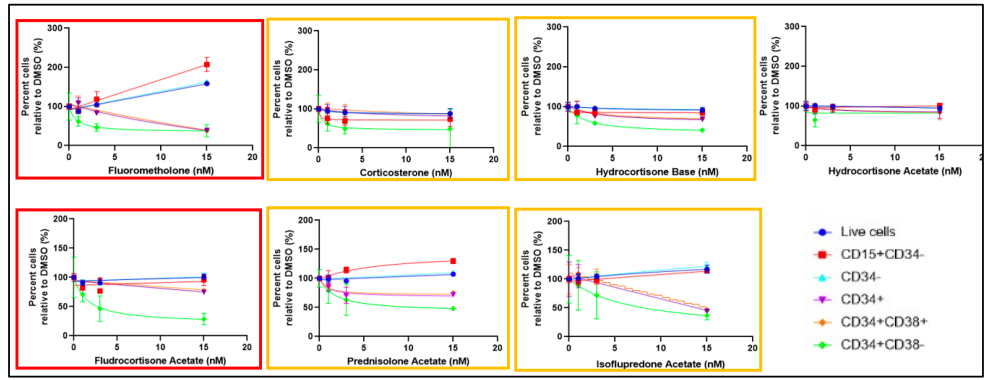

C

**Group B**

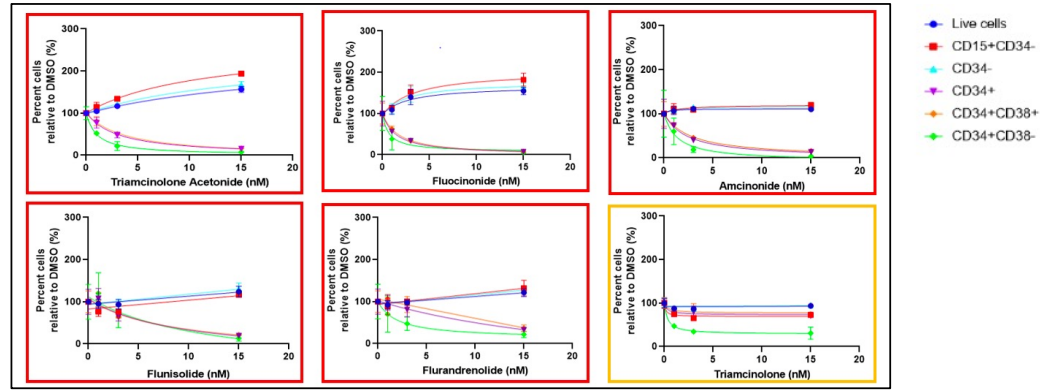

D

**Group C**

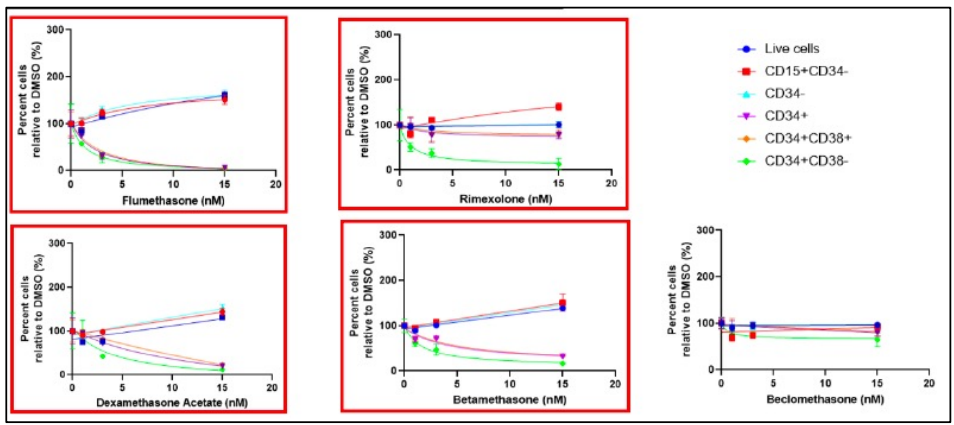

**E**
**Group D1**

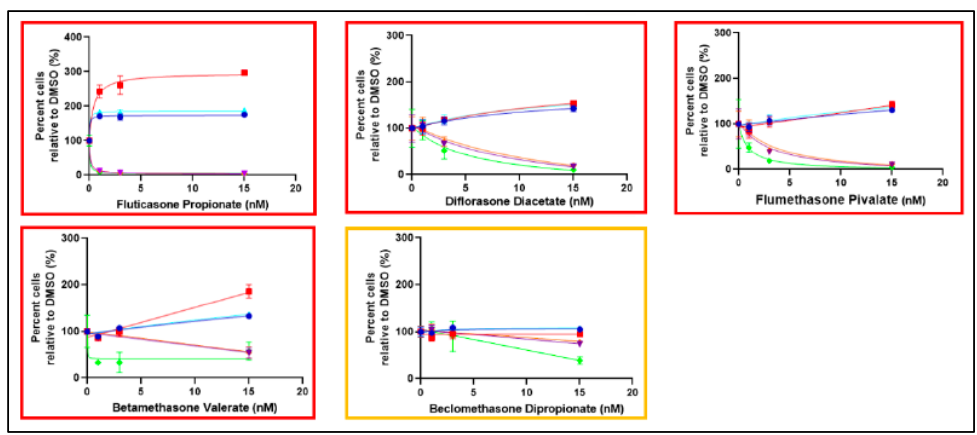

**F**
**Group D2**

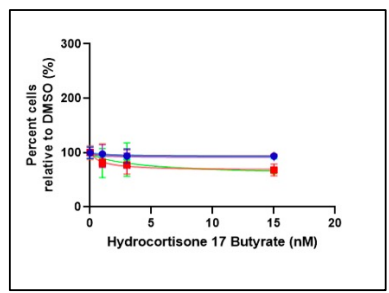

**Supplemental Figure 5: Structural subclass analysis reveals corticosteroids with potent anti-LSC activity and differentiation-inducing capacity.** (A) Dose-response curves for control glucocorticoids MOM and dexamethasone in OCI-AML-8227 cells over six days. Percent cell survival is shown for total live cells, CD15+CD34- (differentiated), CD34- blasts, CD34+CD38+ LPCs, and CD34+CD38- LSC-enriched populations relative to DMSO. Mean ± s.d. (B-F) Twenty-four corticosteroids grouped by Coopman structural classification (Groups A–D2) were screened at 1, 3, and 15 nM. For each compound, cell population viability is plotted relative to DMSO. **Group A (B):** Several compounds showed modest activity; fluorocortisone acetate and fluocinolone acetate displayed moderate LSC depletion and/or differentiation. **Group B (C):**

Triamcinolone acetonide, fluocinonide, amcinonide, and flurandrenolide induced potent LSC depletion and CD34<sup>-</sup> blast expansion, indicating high anti-LSC efficacy. **Group C (D):** Flumethasone, rimexolone, dexamethasone acetate, and betamethasone also reduced LSCs and promoted differentiation. **Group D1 (E):** Fluticasone propionate, diflorasone diacetate, and flumethasone pivalate showed strong LSC depletion with accompanying CD34<sup>-</sup> expansion. **Group D2 (F):** Hydrocortisone 17-butyrate had minimal activity across all populations. Red boxes highlight compounds with strong anti-LSC activity (CD34<sup>+</sup>CD380 LSC-enriched population depletion and CD34<sup>-</sup> blast expansion); orange boxes indicate partial responders; black boxes denote minimal or no response. Line colors correspond to cell populations (legend in panels). Data represent mean ± s.d. Representative one of $n = 2$ biological replicates.

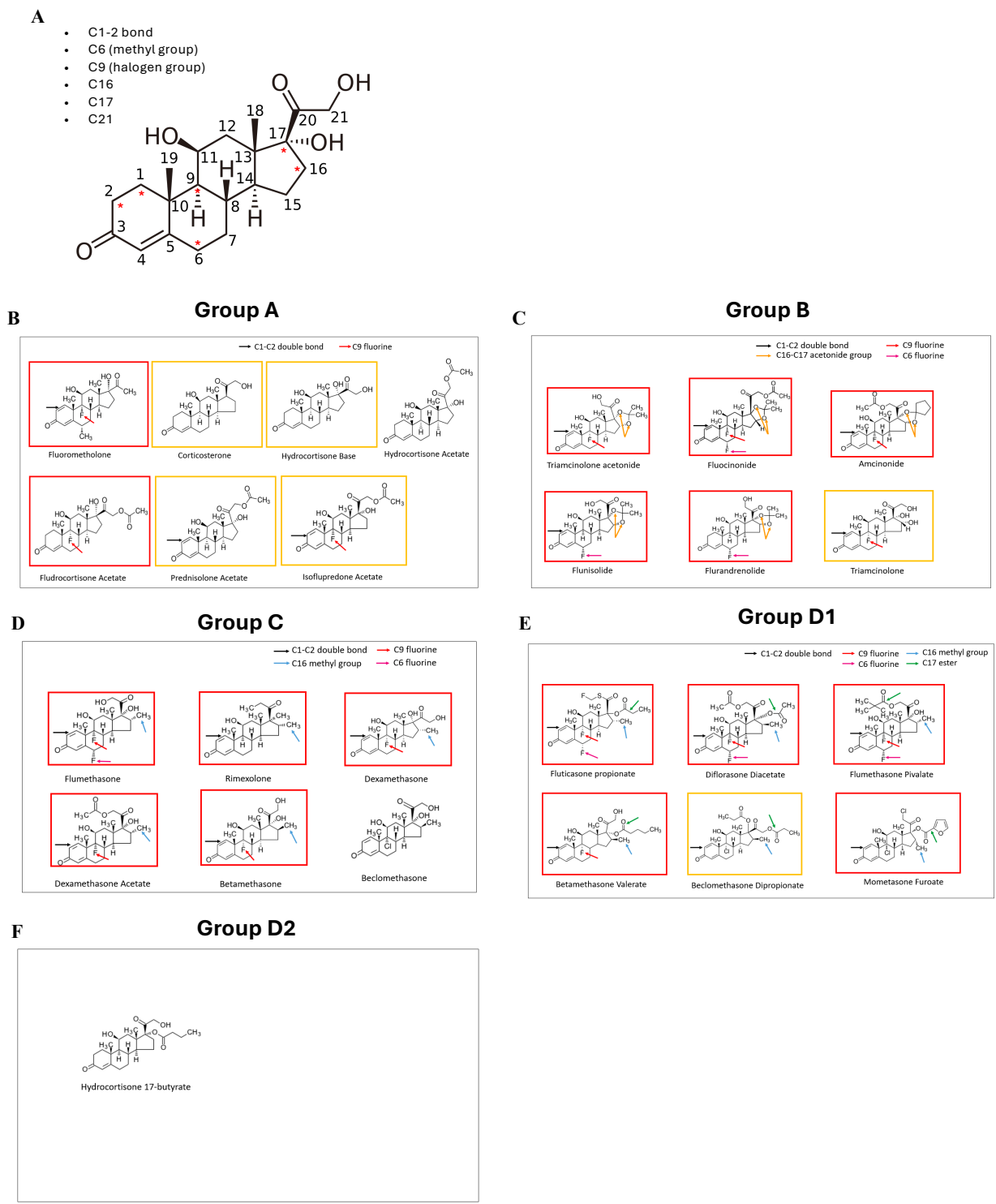

**Supplemental Figure 6: Structural activity relationship identifies features associated with anti-LSC potency among 24 corticosteroids. (A) Reference diagram of the corticosteroid core**

structure, highlighting key modification sites: C1–C2 double bond, C6 methyl group, C9 halogen group, C16/C17 substitutions (e.g., esters), and C21 hydroxyl group. These functional groups vary across corticosteroids and influence glucocorticoid receptor (GR) binding affinity and anti-LSC potency. **(B-F)** Chemical structures of the 24 screened corticosteroids, grouped by Coopman classification (Groups A–D2), annotated to show key activating substitutions. **Group A (B):** Inactive or partially active compounds often lack C9 fluorine or C1–C2 double bonds. Fluorocortisone acetate and fluocinolone acetate possess C9 fluorine and exhibit some activity. **Group B (C):** Active compounds such as triamcinolone acetonide, fluocinonide, and flurandrenolide contain multiple activating features—C9 fluorine, C16/C17 ester substitutions, and/or C1–C2 double bonds. **Group C (D):** Most active compounds (e.g., dexamethasone acetate, flumethasone) display combined C9 halogenation, C16 methylation, and C1–C2 unsaturation. **Group D1 (E):** Fluticasone propionate and diflorasone diacetate show strong anti-LSC activity and contain bulky C17 esters, C6/C9 fluorine, and/or C16 methyl groups. MOM furoate (far right) combines several of these features and was among the most potent hits. **Group D2 (F):** Hydrocortisone 17-butyrate lacks key activating features and showed minimal activity. Red boxes indicate high anti-LSC activity (based on Figure S6); orange boxes indicate partial activity. Structural annotations highlight features correlated with receptor engagement and functional potency.

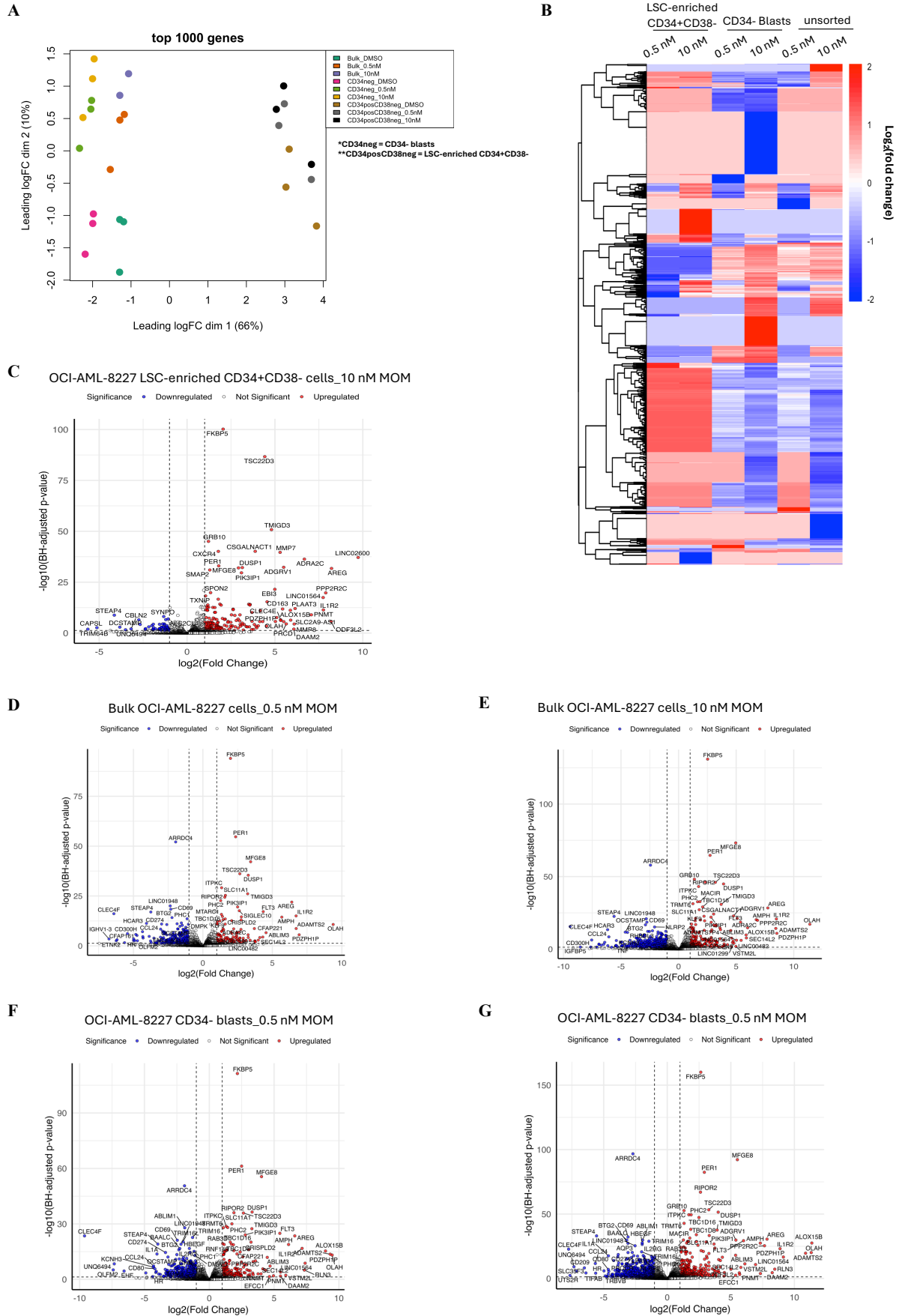

**H**

0.5 nM\_upregulated\_overlap

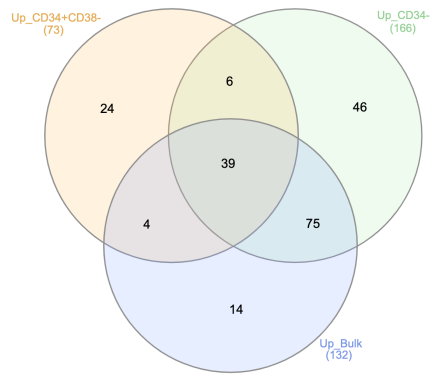

**I**

10 nM\_upregulated\_overlap

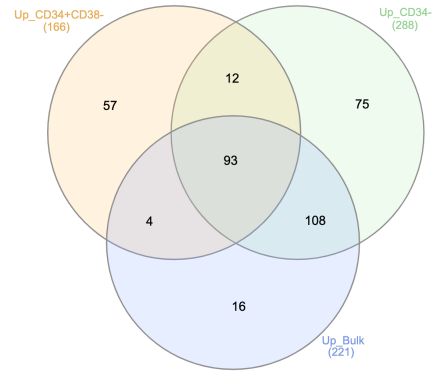

**J**

0.5 nM\_downregulated\_overlap

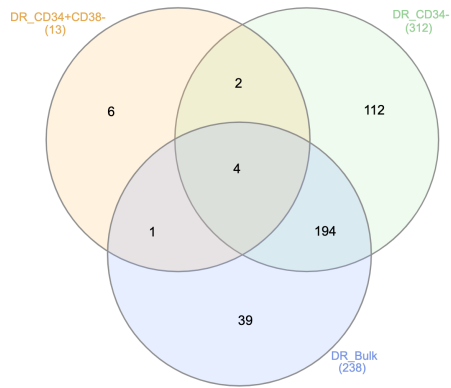

**K**

10 nM\_downregulated\_overlap

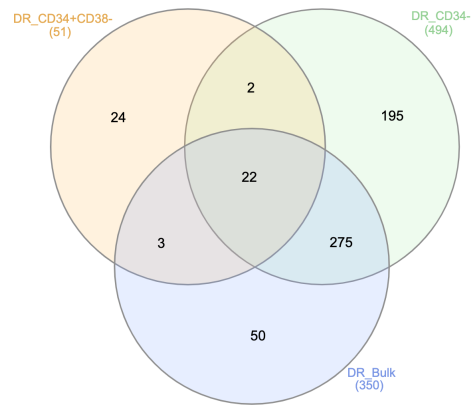

**L**

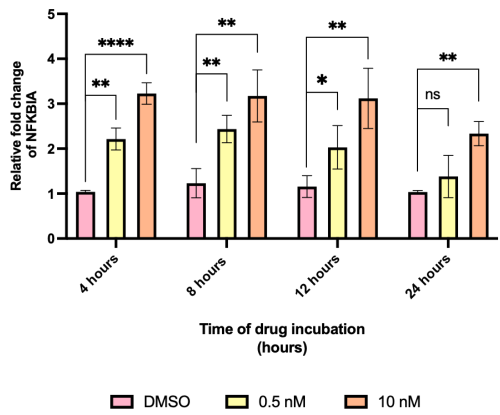

**M**

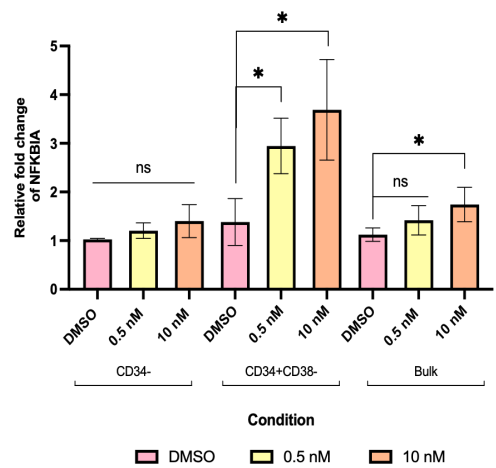

**Supplemental Figure 7: MOM induces early (12 hrs) transcriptional reprogramming in LSC-enriched and bulk OCI-AML-8227 populations as shown by bulk RNA-seq. (A)** PCA using the top 1,000 DEGs highlights distinct transcriptional trajectories induced by MOM across different sorted and unsorted cell populations of OCI-AML-8227. Unsorted OCI-AML-8227 cluster with FACS-sorted CD34<sup>-</sup> blasts, whereas FACS-sorted CD34<sup>+</sup>CD38<sup>-</sup> LSC-enriched population clusters alone, indicating sorting was successful populations respond differently to MOM. Representative of $n = 3$ biological replicates. **(B)** Heatmap of log<sub>2</sub>FC for the top 1,000 variable genes across FACS-sorted CD34<sup>+</sup>CD38<sup>-</sup> LSC-enriched and CD34<sup>-</sup> blast populations, and unsorted populations in OCI-AML-8227 treated with 0.5 or 10 nM MOM. Representative of $n = 3$ biological replicates. **(C-G)** Volcano plots showing differentially expressed genes in bulk/unsorted, FACS-sorted CD34<sup>+</sup>CD38<sup>-</sup> LSC-enriched and CD34<sup>-</sup> blast populations after 12-hour treatment with 0.5 nM or 10 nM MOM, compared to DMSO. Genes with $|\log_2FC| > 1$ and BH-adjusted $p$-value $< 0.05$ are colored red (upregulated) or blue (downregulated). Representative of $n = 3$ biological replicates. **(H-K)** Venn diagrams showing overlap in significantly upregulated or downregulated genes ($FDR < 0.05$, $|\log_2FC| > 1$) between unsorted and FACS-sorted CD34<sup>+</sup>CD38<sup>-</sup> LSC-enriched and CD34<sup>-</sup> blast populations after 12-hour treatment with 0.5 nM **(H, J)** or 10 nM **(I, K)** of MOM. **(L)** Time course of *NFKB1A* transcript levels by RT-qPCR in bulk OCI-AML-8227 cells treated with DMSO, 0.5 nM, or 10 nM MOM for 4, 8, 12, and 24 hours. Representative one of $n = 2$ biological replicates. **(M)** Relative fold change of *NFKB1A* transcript levels at 12 hours in unsorted/bulk and FACS-sorted CD34<sup>+</sup>CD38<sup>-</sup> LSC-enriched and CD34<sup>-</sup> blast populations treated with DMSO or MOM (0.5 nM or 10 nM), showing dose- and population-specific induction. Data are mean ± SD. \*$p < 0.05$, \*\*$p < 0.01$, \*\*\*$p < 0.001$, \*\*\*\*$p < 0.0001$, unpaired t-test. Representative of $n = 1$ biological replicate.

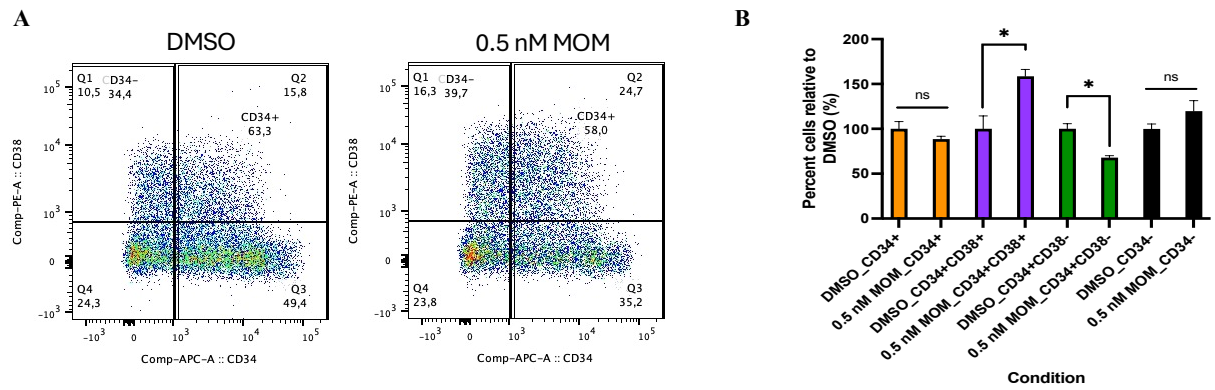

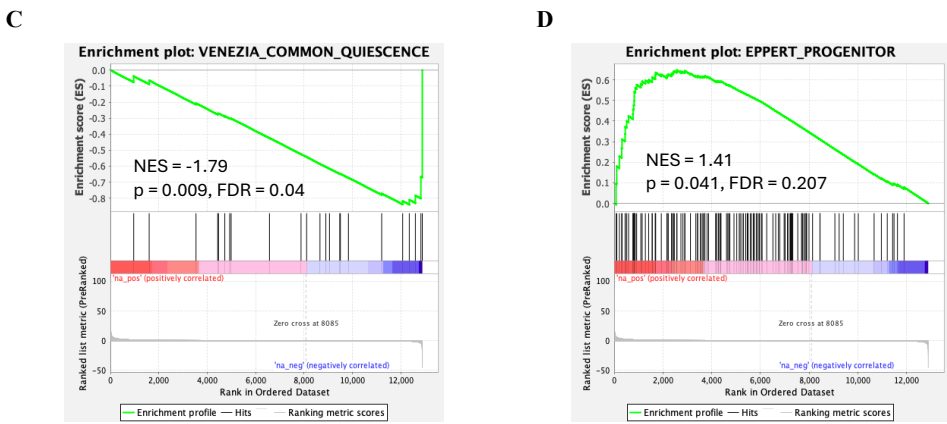

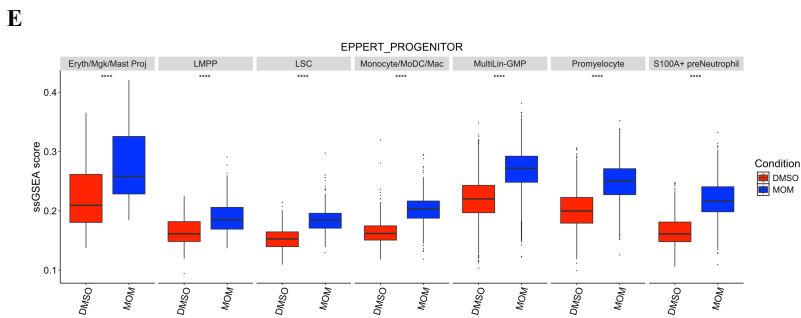

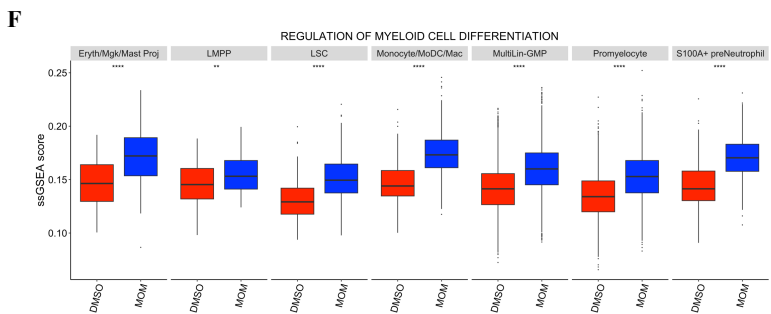

**Supplemental Figure 8: Single cell RNA-seq reveals glucocorticoid-induced loss of quiescence and induction of differentiation signatures in LSC cluster. (A)** Flow cytometry plots of CD34 and CD38 expression of CD34<sup>+</sup> LSPC OCI-AML-8227 cells treated with DMSO or 0.5 nM MOM for 24 h. CD34<sup>+</sup> LSPCs were enriched from OCI-AML-8227 with a purity of 86.5%. MOM reduced CD34<sup>+</sup>CD38<sup>-</sup> LSC-enriched populations, indicating exit from quiescence and primed for differentiation. Representative of $n = 1$ biological replicate. **(B)** Quantification of viable OCI-AML-8227 subpopulations normalized to DMSO control 24 h post-treatment sent for scRNA-seq. A modest decrease in CD34<sup>+</sup>CD38<sup>-</sup> LSC-enriched population and corresponding increase in CD34<sup>+</sup>CD38<sup>+</sup> LPCs and CD34<sup>-</sup> blasts is observed. Unpaired t-test, \* $p < 0.05$, \*\* $p < 0.01$, \*\*\* $p < 0.001$ and \*\*\*\* $p < 0.0001$. Mean ± s.d. Representative one $n = 1$ biological replicate. **(C-D)** GSEA of the LSC cluster (CD34<sup>+</sup>CD38<sup>-</sup>) from 24 h scRNA-seq shows significant negative enrichment of VENEZIA\_COMMON QUIESCENCE and positive enrichment of EPPERT\_PROGENITOR signatures, consistent with differentiation. NES is the normalized enrichment score. Signatures that were significant have normalized p-value $< 0.05$ and FDR $< 0.25$. **(E-F)** ssGSEA shows increased enrichment of gene sets associated with hematopoietic differentiation in the LSC cluster, including **(E)** EPPERT\_PROGENITOR and **(F)** REGULATION\_OF\_MYLOID\_CELL\_DIFFERENTIATION. Wilcoxon test: \*$p < 0.05$, \*\*$p < 0.01$, \*\*\*$p < 0.001$, \*\*\*\*$p < 0.0001$. Representative of $n = 1$ biological replicate.
