## Supplementary Table Titles and Legends for "Glucocorticoids reprogram human AML leukemic stem cells to promote elimination through differentiation and apoptosis"

**Supplemental Table titles**

**Supplemental Table titles**

**Supplemental Table S1: AML patient sample OCI1 cohort (*n* = 15).** This table provides a summary of the OCI1 AML patient samples: primary patient number ID, sample type, %CD34+ and CD34+CD38- cells, sex, FAB subtype, risk group classification, secondary AML status, cytogenetics, initial treatment, and mutations present. N/A: not established.

**Supplemental Table S2: AML patient sample BCLQ cohort (*n* = 9).** This table provides a summary of the BCLQ patient samples: primary AML ID, classification, tissue type, sampling status, age at diagnosis, gender, FAB subtype, blasts percentage, white blood cell count, karyotype, cytogenetic group, AML type, mutations present and LC_50_ to MOM. NA: Not established.

**Supplemental Table S3: AML patient sample OCI2 cohort (*n* = 6).** This table provides information on the AML patient samples from the OCI2: sample ID, Relapse or diagnosis status, FAB subtype, sex, karyotype, number of LSCs in corresponding fractions (CD34+CD38-, CD34+CD38+, CD34-CD38+ and CD34-CD38-), LC_50_ for each fraction for MOM and dexamethasone and mutations present for FLT3-ITD and NPM1. ND: not determined.

**Supplemental Table S4: Response types of AML LSCs to MOM in screened cohorts.** This table summarizes the type of responses seen with MOM treatment in all patient samples tested (*n* = 32) from all 3 cohorts, cell lines and OCI-AML-8227. Table provides the response in LSC fraction, sample ID, mutation information and risk group.

**Supplemental Table S5: Next generation sequencing (NGS) of AML patient samples prior to treatment.** This table includes the mutational profiles of primary AML patient samples that had DNA sequenced. Contains sample ID, mutations present, number of samples with that mutation and type of response, fisher’s test p-value and odd’s ratio.

**Supplemental Table S6: GSEA of RNA-sequencing data from primary AML samples obtained prior to treatment and classified as apoptotic responders.** This table presents gene set enrichment analysis (GSEA) results for OCI-AML-8227 samples classified as apoptotic responders based on post-treatment phenotypes. GSEA was performed on bulk RNA-sequencing data collected **prior to treatment**, using ranked gene expression profiles to assess baseline transcriptional programs. Enrichment was evaluated using hallmark, GO biological process (GOBP), and Reactome gene sets. Reported values include normalized enrichment scores (NES), nominal p-values, and FDR q-values (significance: FDR < 0.25, p < 0.05). Enriched pathways reflect baseline priming toward apoptotic responses.

**Supplemental Table S7: GSEA of RNA-sequencing data from primary AML samples obtained prior to treatment and classified as differentiation responders.** This table contains GSEA results for OCI-AML-8227 samples identified as differentiation responders after treatment, using **pre-treatment bulk RNA-seq data**. Genes were ranked based on baseline expression differences between samples that later underwent differentiation versus those that did not. Enrichment was assessed using hallmark, GOBP, and Reactome databases, with NES, nominal p-values, and FDR q-values reported (significance: FDR < 0.25, p < 0.05). Signatures enriched in this group indicate a predisposition toward differentiation-associated transcriptional programs.

**Supplemental Table S8:** **Structural grouping (Groups A-D2) of 24 glucocorticoids screened in OCI-AML-8227 cells for six days.** List of 24 glucocorticoids tested and the groups they are classified to (Group A, B, C, D1 or D2).

**Supplemental Table S9: Relationship between structural-functional analysis and anti-LSC activity of glucocorticoids.** Provides information on the important groups for anti-LSC activity and number of compounds with these functional groups and those that are effective.

**Supplemental Table S10: Differentially expressed genes (DEGs) identified by RNA-sequencing in bulk OCI-AML-8227 treated with MOM for 12 hours.** Table indicates comparison of 0.5 nM vs DMSO or 10 nM vs DMSO, gene symbols, log_2_CPM, log_2_FC, LR, Pvalue, BH-adjusted p-value, log_2_RPKM for DMSO and corresponding concentrations tested, sign, ranking metric and -log_10_(BH-adjusted p-value).

**Supplemental Table S11:** **Differentially expressed genes (DEGs) identified by RNA-sequencing in FACS-sorted CD34- blasts of OCI-AML-8227 treated with MOM for 12 hours.** Table indicates comparison of 0.5 nM vs DMSO or 10 nM vs DMSO, gene symbols, log_2_CPM, log_2_FC, LR, Pvalue, BH-adjusted p-value, log_2_RPKM for DMSO and corresponding concentrations tested, sign, ranking metric and -log_10_(BH-adjusted p-value).

**Supplemental Table S12: Differentially expressed genes (DEGs) identified by RNA-sequencing in FACS-sorted CD34+CD38- LSC-enriched fraction of OCI-AML-8227 treated with MOM for 12 hours.** Table indicates comparison of 0.5 nM vs DMSO or 10 nM vs DMSO, gene symbols, log_2_CPM, log_2_FC, LR, Pvalue, BH-adjusted p-value, log_2_RPKM for DMSO and corresponding concentrations tested, sign, ranking metric and -log_10_(BH-adjusted p-value).

**Supplemental Table S13: List of overlapping genes identified by RNA-sequencing after 12-hour MOM treatment in bulk (unsorted), FACS-sorted CD34+CD38- LSC-enriched and CD34- blast OCI-AML-8227 cells.** Overlap of genes between CD34- cells, CD34+CD38- LSC-enriched cells and bulk cells at both doses tested (0.5 nM and 10 nM) and for up- and down-regulated genes. Only significant genes are shown. These complement Venn diagram from Supplemental Figure 7H-K.

**Supplemental Table S14: Targeted GSEA of FACS-sorted CD34+CD38- LSC-enriched OCI-AML-8227 cells following 12-hour MOM treatment.** Table shows the list of LSPC signatures and differentiation related signatures used for the targeted analysis. Black box indicates the signatures that were significant (NOM p-value < 0.05 and FDR < 0.25).

**Supplemental Table S15: GSEA results from FACS-sorted CD34+CD38- LSC-enriched OCI-AML-8227 cells treated with MOM for 12 hours.** GSEA of RNA-sequencing ranked lists generated for CD34+CD38- cells. Table provides enriched pathways from KEGG database, gene ontology pathways (GOBP) and hallmark. Pathways with normalized p-value < 0.05 and FDR < 0.25 are significant and shown.

**Supplemental Table S16: GSEA results from FACS-sorted CD34- OCI-AML-8227 blasts treated with MOM for 12 hours.** GSEA of RNA-sequencing ranked lists generated for CD34- blasts. Table provides enriched pathways from KEGG and hallmark databases. Pathways with normalized p-value < 0.05 and FDR < 0.25 are significant and shown.

**Supplemental Table S17: GSEA results from bulk (unsorted) OCI-AML-8227 cells treated with MOM for 12 hours.** GSEA of RNA-sequencing ranked lists generated for bulk cells. Table provides enriched pathways from KEGG database, gene ontology pathways (GOBP) and hallmark. Pathways with normalized p-value < 0.05 and FDR < 0.25 are significant and shown.

**Supplemental Table S18: DEGs identified by scRNA-sequencing across all clusters in OCI-AML-8227 cells following 24-hour MOM treatment.** Table contains gene name, p-value, log_2_FC, FDR, pct.1 and .2, comparison MOM vs DMSO, ranked list by -log_10_pvaluexlog_2_FC and cell_subset pertaining to the 7 clusters identified.

**Supplemental Table S19: GSEA results for the LSC cluster following 24-hour MOM treatment from scRNA-sequencing.** Table contains positively and negatively enriched pathways in LSC cluster from hallmark, reactome and GOBP databases. Black box encapsulates signatures that were significant having normalized p-value < 0.05 and FDR < 0.25.
