## Supplemental Methods for "Glucocorticoids reprogram human AML leukemic stem cells to promote elimination through differentiation and apoptosis"

**Material and Methods**

**Culture and treatment of OCI-AML-8227:**

OCI-AML-8227 cells were cultured in StemSpan™ SFEM II (STEMCELL Technologies) supplemented with 1% penicillin-streptomycin, 10 ng/mL IL-3, IL-6, and G-CSF, 25 ng/mL TPO, 50 ng/mL SCF, and FLT3L (Life Technologies). Cells were treated with glucocorticoids MOM (Tocris), dexamethasone (Sigma), budesonide (Tocris), halcinonide (Selleck), or 0.01–0.02% DMSO, for up to six days. Flow cytometry using a LSR Fortessa with high-throughput sampler (HTS; BD Biosciences) was performed after staining with antibodies to CD34 (APC), CD38 (PE), CD15 (FITC) (Biolegend), and SYTOX^TM^ Blue Dead Stain (Invitrogen^TM^). LC₅₀ values were calculated using GraphPad Prism v10 by nonlinear regression. Time-course assays were conducted at two concentrations per compound (e.g., mometasone at 0.64 and 1.6 nM) and analyzed at 24, 48, 72, and 96 hours.

**Validation in LSC-containing models:**

OP9 stromal cells (ATCC® CRL-2749™) were plated at 5,000 cells/well in Minimal Essential Medium Alpha (α-MEM, 1X) + GlutaMAX™ (-nucleosides) (Fisher Scientific), supplemented with 20% heat-inactivated fetal bovine serum (Wisent), 55 μM β-mercaptoethanol (Fisher Scientific), and 0.01% penicillin-streptomycin. After 48 hours, OP9 media was removed, wells were washed with PBS, and OCI-AML-20 cells (50,000 cells/well) were seeded in IMDM with 10% FBS, 2 mM L-glutamine, 55 μM β-mercaptoethanol, 1% penicillin-streptomycin, and 20 ng/mL GM-CSF (Life Technologies).

MUTZ-3 (DSMZ ACC 295) was cultured in α-MEM (with nucleosides), 20% FBS, 20% 5637-conditioned media, and 1% penicillin-streptomycin. Both models were treated with mometasone and/or dexamethasone for 5–6 days, then stained for CD34 (APC), CD38 (PE), CD15 (FITC), CD14 (AlexaFluor® 700), CD45 (AlexaFluor® 700), Annexin V (Pacific Blue™), and 7-AAD (Biolegend).

**Primary AML samples:**

Samples from the Ontario Cancer Institute (OCI1), Quebec Leukemia Cell Bank, and Nature Medicine cohort (OCI2) were thawed and cultured in OCI-AML-8227 media with 500 nM SR1 and UM729 (STEMCELL Technologies). Glucocorticoids were added for two to four days. Cells were stained with True-Stain Monocyte Blocker™ (Biolegend), and panels as follows:

- **OCI1 cohort:** CD34 (APC-Cy7), CD38 (PE), CD45 (AlexaFluor® 700), CD19 (BV711), CD3 (BV605), CD15 (FITC), CD33 (APC)
- **BCLQ cohort:** same as above but included CD11b (BV650)
- **Nature Medicine cohort (OCI2):** same as above but included CD14 (AlexaFluor® 700) and CD19 (PerCP-Cy5.5).
- Viability was assessed using Annexin V (Pacific Blue™), 7-AAD (Biolegend), or SYTOX^TM^ Blue Dead Stain (Invitrogen^TM^).

**Corticosteroid screen:**

OCI-AML-8227 cells were used for a corticosteroid screen comprising 24 corticosteroids (provided by Dr. Guy Sauvageau, IRIC, Université de Montréal) at 1, 3, or 15 nM for six days. Flow cytometry was used to assess CD34 (APC), CD38 (PE), CD15 (FITC), and SYTOX^TM^ Blue Dead Stain (Invitrogen^TM^). Dose-response curves were used to determine anti-LSC activity and blast induction using GraphPad Prism v10 by nonlinear regression.

**Assessment of NFKBIA expression with mometasone treatment.**

Bulk or FACS-sorted (CD34+CD38- LSC-enriched and CD34- blasts) OCI-AML-8227 cells were plated in corresponding medias and cytokines as mentioned previously. Cells were treated with different concentrations of mometasone (0.5 nM and 10 nM) (Tocris) or control DMSO (Fisher Scientific) and incubated for 4, 8, 12 and 24 hours. Total RNA was extracted using TRIzol^TM^ reagent (Invitrogen) and quantified using a BioDrop^TM^ μLITE spectrophotometer (Montreal Biotech Lab Equipment Inc.).

For cDNA production, 30 ng of 8227 and 200 ng of A549 lung carcinoma cell line (ATCC® CCL-185^TM^) RNA was reversed transcribed in a 20 μL reaction containing 200 units of SuperScript^TM^ III Reverse Transcriptase (Invitrogen), 500 ng of oligo (dT12-18) primers (Invitrogen), 0.5 mM of dNTP mix (Invitrogen), 1X First-Strand Buffer (New England BioLabs inc.), 5 mM DTT (New England BioLabs inc.) and 40 units of RNaseOUT^TM^ Recombinant RNase inhibitor (Invitrogen) using T100TM thermal cycler (Biorad). A total of 2 μL of cDNA was used for qPCR. The following primers were used^1^: *NFKBIA* forward, 5'-AATGCTCAGGAGCCCTGTAAT-3' and reverse, 5'-CTGTTGACATCAGCCCCA CA-3', Actin B forward, 5'-CCTGTACGCCAACACAGTGC-3' and reverse, 5'-ATACTCCTGCTTGCTGATCC-3'. The *NFKBIA* and *Actin B* genes were amplified using Luna® Universal qPCR Master Mix (New England BioLabs inc.). Following 1 cycle of initial denaturation at 95°C for 60 seconds, 45 cycles of 2-step amplification (denaturation at 95°C for 15 seconds and extension at 60°C for 30 seconds + plate read) and 1 cycle of melting (95°C for 10 seconds, 65°C for 60 seconds and finally 97°C for 1 seconds, continuous) using the Lightcycler 96 (Roche).

The results are presented as Ct values, defined as the threshold PCR cycle number at which an amplified product is first detected. A standard curve was generated (5-fold dilutions) using cDNA from A549 for *NFKBIA* and *Actin B* to assess primer efficiency. To establish the change in expression of *NFKBIA* (gene of interest), the average Ct values were calculated for *NFKBIA* and *Actin B* untreated (DMSO control) and treated samples. The ΔCt was determined as the difference between the mean Ct values for *NFKBIA* minus the mean Ct values of *Actin B*. The 2-ΔΔCt method was used to assess change in expression of *NFKBIA* for treated samples relative to DMSO control. Cells were stained with antibodies to CD34 (APC), CD38 (PE), CD15 (FITC) (Biolegend) and SYTOX^TM^ Blue Dead Stain (Invitrogen^TM^). Flow cytometry was performed using an LSRFortessa fitted with a high-throughput sampler (HTS).

**DNA and RNA extraction**

DNA and RNA was extracted from the OCI1 (*n* = 15) and OCI2 cohort samples (*n* = 6) prior to treatment. 1 million cells were collected, and cell pellets and lysates were prepared by following DNeasy® Blood & Tissue and RNeasy® Mini kits (QIAGEN). RNA isolated from primary samples were sent to Génome Québec for RNA sequencing and were analyzed via Compute Canada (see below for more details). The data was then processed using R and GSEA. Next-Generation DNA sequencing and processing was performed by Dr. Luca Cavallone from Dr. Yury Monczak’s lab, and R was used for performing a Fisher’s test for significance. For apoptosis analysis, samples from OCI2 were not included as they lacked apoptosis information. For differentiation analysis, all samples were included.

For OCI-AML-8227, CD34+CD38− LSC-enriched, CD34− blasts and bulk OCI-AML-8227 cells were sorted by FACS, treated with DMSO (0.02%) or MOM (0.5 or 10 nM) for 12 hours (*n* = 3 biological replicates), and processed for RNA extraction using TRIzol (Invitrogen).

**RNA-sequencing of OCI-AML-8227 (bulk and FACS-sorted) treated with mometasone for 12 hours.**

OCI-AML-8227 cells (*n* =3 biological replicates) were collected and washed twice with 2% CCS (Fisher Scientific) in PBS (Life Technologies) and then counted. Some OCI-AML-8227 cells (~1-2 million) were set aside for the bulk condition (unsorted). Cells were then stained with antibodies to CD34 (APC) and CD38 (PE) (Biolegend) for 30 minutes. Cells were washed with 2% CCS in PBS after staining and filtered using Falcon® round-bottom polystyrene tubes with 35 μm cell strainer cap (VWR). Cells were spun down after filtering and resuspended in a concentration of 20 million cells/mL with 2% CCS in PBS. Cells were sorted using BD FACSAria^TM^ Fusion cell sorter into CD34- and CD34+CD38- populations in SFEM II with 0.01% penicillin-streptomycin. After sort, cells were spun down, counted and plated (25,000 per well) in 96-well flat bottom non-TC treated plates (bulk vs CD34- vs CD34+CD38-). Cells were treated with mometasone at different concentrations (DMSO only 0.02%, 0.5 nM and 10 nM) and incubated for 12 hours at 37°C and 5% CO_2_. After 12 hours of treatment with mometasone, total RNA was extracted using TRIzol^TM^ reagent (Invitrogen) and quantified using a BioDrop^TM^ μLITE spectrophotometer (Montreal Biotech Lab Equipment Inc). RNA samples were submitted to Genome Quebec (Montreal, CA) for sequencing analysis.

Total RNA was quantified using a NanoDrop Spectrophotometer ND-1000 (NanoDrop Technologies, Inc.) and its integrity was assessed on a 2100 Bioanalyzer (Agilent Technologies). RNA samples were submitted to Genome Quebec (Montreal, CA) for sequencing analysis. Libraries were generated from 250 ng of total RNA using the Illumina® Stranded mRNA Prep, Ligation kit (Illumina), as per the manufacturer’s recommendations. Libraries were quantified using the KAPA Library Quantification Kits - Complete kit (Universal) (Kapa Biosystems). Average size fragment was determined using a LabChip GXII (PerkinElmer) instrument. Libraries were sequenced using the NovaSeq 6000 S4 PE100 BP sequencing system. The libraries were normalized and pooled and then denatured in 0.02N NaOH and neutralized using HT1 buffer. The pool was loaded at 175pM on an Illumina NovaSeq S4 lane using Xp protocol as per the manufacturer’s recommendations. The run was performed for 2x100 cycles (paired-end mode). A phiX library was used as a control and mixed with libraries at 1% level. Base calling was performed with RTA v3.4.4. Program bcl2fastq2 v2.20 was then used to demultiplex samples and generate fastq reads. Compute Canada software was used to map the reads to the reference genome and count the read numbers mapped to each gene. Differential expression analysis between two groups or more was performed using EdgeR package, implemented in genpipe’s RNA-seq pipeline (v4.4.2) while taking replicates into account.

Genes with a BH-adjusted p-value of 0.05 and log_2_ fold change > 1 and <-1 were assigned as differently expressed and volcano and box plots were created using R package ggplot2. Gene set enrichment analysis (GSEA) (targeted and non-targeted) with default parameters was used to identify enriched pathways. VennDiagram was used to identify overlap between genes. Cells were also collected and washed with 2% CCS in PBS to stain with antibodies against CD34 (APC), CD38 (PE), CD15 (FITC) (Biolegend) and SYTOX^TM^ Blue Dead Stain (Invitrogen^TM^). Flow cytometry was performed using an LSR Fortessa fitted with a high-throughput sampler (HTS).

**Single cell RNA sequencing.**

OCI-AML-8227 cells were enriched using SFEM II and the EasySep^TM^ Magnet and EasySep™ CD34 Positive Selection Kit II protocol (Catalog #17856, STEMCELL Technologies). A total of four washes were performed instead of five to keep some CD34- blasts in the purified sample. Cells were then washed in SFEM II and counted using a hemocytometer to determine final yield. Some cells were stained with antibodies against CD34 (APC), CD38 (PE), CD15 (FITC) (Biolegend) and SYTOX^TM^ Blue Dead Stain (Invitrogen^TM^). Flow cytometry was performed using a LSR Fortessa to determine purity of CD34+ and CD34- fractions. Cells were then plated (50,000 cells/well) and treated with 0.05% DMSO only (Fischer Scientific) and 0.5 nM of Mometasone (Tocris).

Cells were incubated for 24 hours and then collected, washed and resuspended in PBS with 0.04% BSA (Life Technologies). We aimed to capture ~10,000 cells per sample. An aliquot of cells was used for LIVE/DEAD viability testing (Thermo Fisher Scientific). Single-cell libraries were generated using the 10x Genomics Chromium X instrument and Chromium Next GEM Single Cell 3ʹ GEM, Library & Gel Bead Kit v3.1 (10x Genomics) according to the manufacturer’s protocol. Briefly, cells suspended in reverse transcription reagents, along with gel beads, were segregated into aqueous nanoliter-scale gel bead-inemulsions (GEMs). The GEMs were then reverse transcribed in a T1000 Thermal cycler (Bio-Rad) programed at 53ºC for 45 min, 85ºC for 5 min, and hold at 4ºC. After reverse transcription, single-cell droplets were broken, and the single-strand cDNA was isolated and cleaned with Cleanup Mix containing DynaBeads (Thermo Fisher Scientific). cDNA was then amplified with a T1000 Thermal cycler programed at 98ºC for 3 min, 12 cycles of (98ºC for 15 s, 63ºC for 20 s, 72ºC for 1 min), 72ºC for 1 min, and hold at 4ºC. Subsequently, the amplified cDNA was fragmented, end repaired, A-tailed and index adaptor ligated, with SPRIselect Reagent Kit (Beckman Coulter) with cleanup in between steps. Post-ligation product was amplified with a T1000 Thermal cycler programed at 98ºC for 45 s, 12 cycles of (98ºC for 20 s, 54ºC for 30 s, 22 72ºC for 20 s), 72ºC for 1 min, and hold at 4ºC. The sequencing-ready libraries were cleaned up with SPRIselect, quality controlled for size distribution and yield (LabChip GX Perkin Elmer), and quantified using qPCR (KAPA Biosystems Library Quantification Kit for Illumina platforms).

Libraries were loaded on NovaSeq 6000 and sequenced using the following parameters: 28 bp Read1, 8 bp Index i7, 8 bp Index i5 and 150 bp Read2.

Raw sequencing BCL files from the Illumina sequencing were demultiplexed into paired-end, gzip-compressed FASTQ files using Illumina's bcl2fastq. Following this using cellranger count version 7.1.0 (10X genomics), reads were aligned to the GRCh38 human reference genome, and transcript counts were quantified for each annotated gene within every cell. The resulting UMI count matrices (genes Å~ cells) were then provided as input to Seurat suite.

**Single-cell RNA sequencing analysis.**

Cell Ranger v7.1.0 (10x Genomics) was used to map reads to the human reference transcriptome, GRCh38 v1.2.0 (also from 10x Genomics). Seurat v5.1.0^3^ within R v4.3^4^ for was used for data normalization, integration and visualization. Cells were filtered out if they had: more than 15% mitochondrial gene expression, less than 250 genes identified, less than 500 UMIs as well as, those cell that had a low novelty score (log10 GenesPerUMI < 0.8). Similarly, the described steps by DoubletFinder v2.0.4^5^ were followed to identify and remove doublets from the dataset.

“SCTranform” was used to perform normalization regressing out mitochondrial expression, followed by integration as described by Seurat using the “RPCA” method. The umaps, violin plots, heatmaps and related figures were made using a combination of Seurat v5.1.0 and Tidyverse v2.0.0^6^ tools.

**Cell type prediction using ANNCAST.**

Cell type prediction was performed using the ANNCAST classifier v1.2^7^, an artificial neural network-based model trained specifically for AML cells. To ensure compatibility with the classifier, outdated gene symbols from Ensembl84 were first converted to Ensembl97 by mapping GeneIds to the appropriate gene symbol version used by ANNCAST. Gene expression matrices were then normalized by sequencing depth, scaled to a constant depth of 10,000 UMIs, and log-transformed. To reduce the likelihood of ambiguous cell type assignments, any cell with a confidence score below 0.8 was re-assigned to the majority cell type among its five nearest neighbors (based on PCA embeddings) that had high-confidence scores (≥0.8). Because the ANNCAST classifier assigns cells to one of 18 high-confidence subtypes—and some of these subtypes contained relatively few cells—these were further grouped into seven broader categories: LSC, MultiLin-GMP, LMPP, Promyelocyte, S100A+ preNeutrophil, and Monocyte/MoDC/Mac, Erythrocyte/Megakaryocyte/Mast progenitors were also grouped to obtain subsets of suitable size for differential gene expression analysis. The resulting cell type assignments were reviewed and validated using a panel of key cell lineage marker genes.

**Differential gene expression (DGE).**

Differential gene expression (DGE) analysis was performed to assess the impact of treatment on the transcriptional profile. Each treatment condition was compared to the DMSO control. Comparisons were conducted within unsupervised clusters, within ANNCAST-categorized cell types, and within cell types stratified by tumor subclone identity. A zero-inflated model was applied to the log-normalized gene expression data using the MAST v1.28 statistical framework^7^. The number of detected genes per cell was included as a covariate alongside the treatment effect. P-values were adjusted for multiple testing using the Benjamini-Hochberg (FDR) method. For the DGE analysis within tumor subclones, only clone–cell type combinations with at least 100 cells were included in the analysis to ensure robust differential expression testing.
